# Tfap2a is a novel gatekeeper of differentiation in renal progenitors during kidney development

**DOI:** 10.1101/460105

**Authors:** Brooke E. Chambers, Gary F. Gerlach, Karen H. Chen, Eleanor G. Clark, Ignaty Leshchiner, Wolfram Goessling, Rebecca A. Wingert

## Abstract

Renal functional units known as nephrons undergo patterning events during development that create a segmental array of cellular populations with discrete physiological tasks. Knowledge about the terminal differentiation programs of each nephron segment has central importance for understanding kidney disease and to advance regenerative medicine, as mammalian nephrons grown in organoid cultures from pluripotent cells fail to terminally differentiate. Here, from a novel forward genetic screen using zebrafish we report the discovery that *transcription factor AP-2 alpha* (*tfap2a)* coordinates a gene regulatory network that controls the progression of nephron distal segment progenitors into the differentiated state. Overexpression of *tfap2a* rescued differentiation in mutants and caused ectopic expression of distal segment markers in wild-type nephrons, indicating *tfap2a* is sufficient to instigate the distal segment differentiation program. *tfap2a/2b* deficiency exacerbated distal nephron segment differentiation defects, revealing functional redundancy where *tfap2a* has a dominant role upstream of its family member. With further genetic studies, we assembled a blueprint of the *tfap2a* gene regulatory network during nephrogenesis. We demonstrate that *tfap2a* acts downstream of *Iroquois homeobox 3b*, a conserved distal lineage transcription factor. *tfap2a* controls a circuit consisting of *irx1a, tfap2b,* and genes encoding solute transporters that dictate the specialized metabolic functions of the distal nephron segments, and we show for the first time that this regulatory node is distinct from the pathway circuits controlling aspects such as apical-basal polarity and ciliogenesis during the differentiation process. Thus, our studies reveal new insights into the genetic control of differentiation, where *tfap2a* regulates the suite of segment transporter traits. These findings have relevance for understanding renal birth defects, as well as efforts to recapitulate nephrogenesis *in vivo* to make functional units that can facilitate organoid applications such as drug discovery and regenerative therapies.

**Summary Statement:** Here, we report for the first time that *transcription factor AP-2 alpha* (*tfap2a*) controls the progression from nephron progenitor into the fully differentiated state. This fundamentally deepens our knowledge about the genetic control of kidney development.

## Introduction

Vertebrate kidney ontogeny involves the reiterative formation and degradation of several structures from the intermediate mesoderm: the pronephros, the mesonephros, and the metanephros (Saxen, 1987). In amniotes, the metanephros serves as the final kidney form, while in lower vertebrates, such as fish and frogs, the mesonephros functions as the adult kidney. Importantly, all the kidney versions are comprised of conserved functional units called nephrons (Dressler, 2006). The nephron is comprised of a blood filter, a segmented epithelial tubule, and a collecting duct. Each of these anatomical nephron parts modifies the filtrate in a stepwise fashion to perform the vital tasks of excretion, pH balance, and fluid homeostasis. Occurring in approximately 1 in 500 births, Congenital Anomalies of the Kidney and Urinary Tract (CAKUT) are among the most common birth defects and are the primary cause of pediatric end stage renal disease (ESRD) (Airik and Kispert, 2007; Song and Yosypiv, 2011). The shared etiology across these diverse conditions is the aberrant development of nephrons stemming from genetic dysregulation (Schedl, 2007). To this end, it is imperative to understand the signals that coordinate nephron formation during renal organogenesis.

The zebrafish pronephros has emerged as a genetically tractable vertebrate model to study the molecular mechanisms regulating nephron segment development events (Wingert et al., 2007; Wingert and Davidson, 2008; Wingert and Davidson, 2011). The embryonic zebrafish is transparent in nature, and its pronephric kidney is structurally simple, consisting of two bilateral nephrons that make it an excellent model to study renal progenitor changes *in vivo* (Naylor et al., 2017). Like other vertebrate nephrons, the zebrafish pronephros is patterned into distinct proximal and distal epithelial segments (Wingert et al., 2007; Wingert and Davidson, 2008). Further, zebrafish mirror fundamental processes of mammalian nephron formation such as the mesenchymal to epithelial transition (MET) of renal progenitors, establishment of apical-basal polarity, lumen formation, ciliogenesis, and formation of specialized segment populations (Gerlach and Wingert, 2013). While there has been significant progress in understanding nephron segment patterning in recent years (Desgrange and Cereghini, 2015; Lindström et al., 2015; Chung et al., 2017), the pathways that dictate segmental terminal differentiation are far from understood.

Transcription factors play a central role in operating the genetic networks that orchestrate renal cell fate acquisition and nephron segment patterning (Desgrange and Cereghini, 2015; Lindström et al., 2015). Advances in single-cell RNA sequencing and gene expression analysis in both the embryonic murine and human kidneys have brought to light an inventory of factors mapped to distinct regions of developing nephrons (Lindström et al., 2018a, b). Although these studies have provided a detailed transcription factor localization atlas that is time-dependent, the functions of these factors in nephrogenesis have not yet been fully elucidated. One uncharacterized gene is *Transcription Factor AP-2 Alpha (TFAP2A)*, which clustered with developing medial/distal tubule signatures (Lindström et al., 2018b). *TFAP2A* is a member of the AP-2 transcription factor family (AP-2α, AP-2β, AP-2γ, AP-2δ and AP-2∊), whose proteins share highly conserved dimerization and DNA binding motifs across vertebrates (Fig. S1). AP-2 factors bind to GC-rich promoter sequences, and can homodimerize and heterodimerize with one another (Eckert et al., 2005). During development, these factors have been shown to exercise redundant and unique functions depending on the tissue context (Eckert et al., 2005).

*Tfap2a* and family member *Tfap2b* also have overlapping expression patterns during vertebrate embryogenesis in neural crest derivatives, surface ectoderm, and the kidney (Moser et al., 1997; Knight et al., 2003; Knight et al., 2005). Surprisingly, the elimination of *Tfap2a* and *Tfap2b* in mice results in completely different phenotypic outcomes. *Tfap2a* knockout mice die perinatally and display a suite of pleiotropic features that include craniofacial alterations, incomplete neural tube closure, hypoplastic hearts and kidneys (Schorle et al., 1996; Zhang et al., 1996; Brewer and Williams, 2004; Brewer et al., 2004). In contrast, *Tfap2b* null mice exhibit patent ductus arteriosus and die shortly after birth due to acute renal failure with elevated apoptosis (Moser et al., 1997; Hilger-Eversheim et al., 2000; Wang et al., 2018). Because *Tfap2b* mutants exhibit less severe phenotypes, this factor is proposed to share redundant functions with *Tfap2a* during development (Eckert et al., 2005; Kerber et al., 2001). An example in support of this relationship is that *Tfap2a* plays a more dominant role than *Tfap2b* in the development of branchial arches in mice (Van Otterloo et al., 2018).

Genetic defects in the AP-2 factors are associated with several human diseases. Autosomal dominant *TFAP2A* mutations in humans cause branchio-oculo-facial syndrome (BOFS), which primarily affects craniofacial tissue (Milunsky et al., 2008). Additionally, human *TFAP2A* lesions are associated with multicystic dysplastic kidney defects, but the mechanisms have remained unexplored. Dominant-negative mutations in human *TFAP2B* cause Char Syndrome, which affects heart, face, and limb development (Satoda et al., 2000). Despite the previously documented renal phenotypes associated with *Tfap2a* and *Tfap2b* deficiency in rodents, these factors have not been studied further in the context of kidney development. For example, nephron segmentation has not been analyzed in either *Tfap2a* or *Tfap2b-* deficient murine models. Nevertheless, *Tfap2a/tfap2a* has been extensively studied in the vertebrate neural crest, where it facilitates specification and differentiation through a complex genetic regulatory network (Knight et al., 2003; Knight et al., 2005; Holzschuh et al., 2003; Barrallo-Gimeno et al., 2003; O’Brien et al., 2004; Li and Cornell, 2007; Hoffman et al., 2007; Van Otterloo et al., 2010; de Crozé et al., 2011; Wang et al., 2011; Bhat et al., 2012; Green et al., 2014; Kantarci et al., 2015; Seberg et al., 2017). These studies provide a valuable framework with which to consider the roles of *Tfap2a/tfap2a* in other tissues, where it is likely to also mediate genetic networks.

Here, we report the novel zebrafish nephron segment mutant, *terminus* (*trm)*, which was isolated in a forward haploid genetic screen. Employing whole genome sequencing, we identified a mutation that blocks proper splicing of *tfap2a*. While *tfap2a* deficient nephrons have normal distal segment pattern formation, display normal epithelial polarity and cilia development, they experience a block in other aspects of terminal differentiation, resulting in the loss of solute transporter expression within distal segments. Interestingly, *tfap2a* is sufficient to induce ectopic expression of distal segment markers in adjacent segment domains. We found that *tfap2b* functions redundantly and downstream of *tfap2a* to turn on the distal nephron solute transporter program. Further, *tfap2a* articulates with the Iroquois homeobox transcription factors *irx1a* and *irx3b*, which are regulators of intermediate/distal nephron identity. Our study reveals for the first time that *tfap2a* controls a gene regulatory network that serves as a gatekeeper of terminal differentiation during nephron segment development, and establishes a new paradigm that will be valuable to deepen our knowledge of cell differentiation mechanisms in the kidney.

## Results

### Forward genetic screen identifies *tfap2a* as a novel regulator of nephron development

There remain many gaps in our understanding of the genetic blueprint needed to orchestrate renal stem cell fate decisions and nephron segment formation during kidney ontogeny. The embryonic zebrafish kidney, or pronephros, is a practical genetic model for nephron segmentation (Gerlach and Wingert, 2013). At 24 hours post fertilization (hpf), the pronephros is fully formed and exhibits a very simple organization consisting of two parallel nephrons (Fig. 1A), making cellular changes easy to detect (Poureetezadi and Wingert, 2016). Each nephron is comprised of a blood filter, a series of proximal and distal segments that reabsorb and secrete molecules, and a collecting duct to transport waste (Fig. 1A) (Wingert et al., 2007).

**Figure 1:**
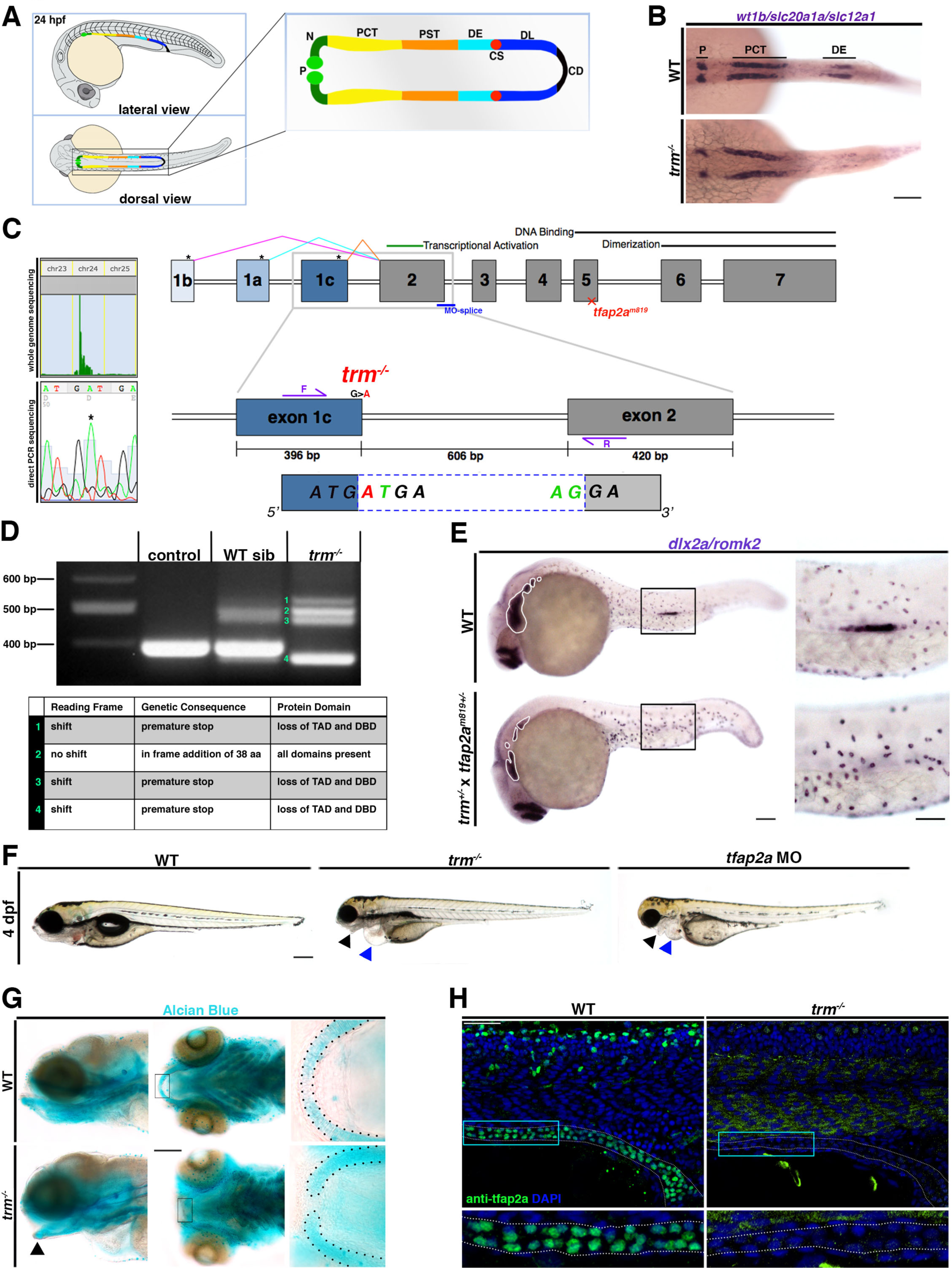
Forward genetic screen reveals *tfap2a* is necessary for nephrogenesis in the developing zebrafish pronephros. **A.** Schematic depicts lateral and dorsal views of fully segmented pronephros in 24 hpf zebrafish embryo (P: podocytes, N: neck, PCT: proximal convoluted tubule, DE: distal early segment, CS: corpuscle of Stannius, DL: distal late segment, CD: collecting duct). **B.** Screening approach by WISH of alternating nephron compartment markers *wt1b/slc20a1a/slc12a1* (P/PCT/DE) in WT and *trm* at 24 hpf. Scale bar = 70µm. **C.** SNPtrack results from whole genome sequencing concentration of SNPs at chromosome 24. Confirmation of G -> A *tfap2a* mutation by direct PCR sequencing of *trm*^-/-^ embryos. Exon diagram of *tfap2a* depicts 3 alternative spliceoforms (pink, cyan, orange lines). Black *’s indicate alternative start sites. *tfap2a* MO-splice (blue) targets 3’ end of exon 2. *tfap2a^m819^* lesion (red x) generates stop codon in exon 5. *trm* G > A mutation (red) maps to 3’ end of exon 1c. Green letters indicate conserved splice residues. RT-PCR primers used for RT-PCR analysis flank intron 1-2 (purple arrows). **D.** RT-PCR analysis of control (WT) embryos, WT *trm* siblings, and *trm*^-/-^. Mutant bands are labeled 1-4 in green. Table indicates predicted genetic consequence from sequencing the mutant bands. (TAD:transcriptional activation domain, DBD:DNA binding domain**). E.** Failure to complement revealed by WISH analysis of *dlx2a* (pharyngeal arches outlined in white) and *romk2* (DE) in WT and *trm^+/^-* x *tfap2a*^m819+/-^ compound mutants. Scale bars = 70 µm, 35 µm. F. Live imaging at 4 dpf reveals abnormal craniofacial cartilage (black arrowhead) and pericardial edema (blue arrowhead) in *trm*^-/-^ and *tfap2a* morphants. Scale bar = 200 µm. **G.** Alcian Blue cartilage staining in WT and *trm*^-/-^ at 4 dpf. Gaping jaw phenotype indicated by black arrowhead. Black dotted lines are utilized to trace Meckel’s cartilage. Scale bar = 100 µm. **H.** Whole mount IF of Tfap2a protein in WT and *trm*^-/-^ at 24 hpf. White dotted lines delineate pronephric tubule. Cyan box denotes 40x optical zoom. Scale bar = 30 µm.

To identify novel renal regulators, we performed a forward genetic haploid screen in zebrafish. We obtained maternal gametes from the F1 generation, applied ultraviolet light inactivated sperm to generate F2 haploid embryos, and assayed them for nephron segment defects by whole mount *in situ* hybridization (WISH) (Kroeger et al., 2014). For our assay we applied a mixture of probes that specifically localize to alternating compartments of the pronephros: podocytes (*wt1b*), proximal convoluted tubule (PCT) (*slc20a1a*), and the distal early segment (DE) (*slc12a1*) (Fig. 1B). Through this multiplex assay we isolated the nephron mutant *terminus* (*trm)* which affected DE segment development based on abrogated *slc12a1* expression within the pronephros (Fig. 1B).

We performed whole genome sequencing to identify the causative gene for the *trm* phenotype (Leshchiner et al., 2012). Using SNP track software analysis, the location of the genetic lesion was mapped to chromosome 24 (Fig. 1C). We used previously described thresholds to enrich results (Ryan et al., 2013) and discovered the gene *tfap2a* was a high scoring candidate at the chromosome 24 locus, where there was a G -> A substitution that was predicted to disrupt splicing at the splice donor site of exon 1. We performed direct PCR sequencing on *trm* mutants and wild-type (WT) siblings, and confirmed this genetic change (Fig. 1C). To characterize how this mutation affected splicing, we conducted transcript analysis. RT-PCR on total RNA isolated from *trm* mutants revealed four aberrant *tfap2a* spliceoforms compared to WTs (Fig. 1D). One aberrant transcript encoded an in-frame addition of 38 amino acids (Fig. 1D), which may possess native function or have dysfunctions associated with protein folding or stability. The other three transcripts encoded premature stop codons (Fig. 1D). These aberrant *trm* transcripts are predicted to truncate the essential transcriptional activation and DNA binding domains in the Tfap2a protein (Fig. 1D).

Next, we explored whether the loss of *tfap2a* function in *trm* mutants was the sole origin of their renal phenotype. To do this, we performed complementation tests between *trm* and *tfap2a^m819^,* the latter which encodes a nonsense allele, followed by phenotype assessment with WISH and finally genotyping analysis. Compound *trm*^+/-^;*tfap2a*^m819+/-^ heterozygote embryos displayed the loss of *romk2* expression within the pronephros DE segment and reduced *dlx2* expression in the neural crest as well (Fig. 1E). This result indicates that the alleles failed to complement one another, which most likely indicates that the same gene is affected, and provided powerful evidence that disruption of *tfap2a* expression alone underlies the *trm* phenotype.

Until now, *tfap2a* has been known as essential for neural crest and epidermis differentiation. Thus, we assessed whether *trm* mutants evinced hallmarks of *tfap2a* deficiency. With analysis of live morphology at 4 dpf, we found that *trm* developed abnormal craniofacial cartilage and pericardial edema, which was phenocopied upon *tfap2a* morpholino knockdown (Fig. 1F). RT-PCR analysis confirmed that this *tfap2a* MO effectively disrupts splicing (Fig. S2). We examined facial cartilage using Alcian blue staining, where we found *trm* possesses defects in Meckel’s cartilage and pharyngeal arch structures (Fig. 1G). The cartilage phenotypes observed in *trm* are consistent with the documented neural crest *tfap2a* mutant alleles *lockjaw* and *mont blanc* (Knight et al., 2003; Barrallo-Gimeno et al., 2003). *trm* also displayed disrupted craniofacial vasculature formation as indicated by o-dianisidine staining (Fig. S4). Further, when we assayed for *tfap2a* protein by whole mount immunofluorescence (IF), we detected no pronephric expression in *trm* mutants as compared to WT at 24 hpf. The absence of *tfap2a* protein expression in the mutant pronephros indicates that the ultimate consequence of *trm* allele is a bona-fide loss-of-function. In light of the mutation site, these IF data further indicate that the *tfap2a* exon1c spliceoform encodes the dominant protein variant that is active during kidney development. In sum, these results show that *trm* mutants exhibit many features of *tfap2a* deficiency, and reveal for the first time that *tfap2a* is needed for nephrogenesis—specifically for proper emergence of the DE segment population.

### *tfap2a* and *tfap2b* are coexpressed dynamically during nephron development and function redundantly to induce regimens of distal segment solute transporter genes

Of the AP-2 family of transcription factors, only *tfap2a* and *tfap2b* have been reported as expressed in the developing zebrafish pronephros (Knight et al., 2005; Sugano et al., 2017). Zebrafish *tfap2a* and *tfap2b* genes are closely related, as they share overall 65% amino acid sequence identity, with highly similar DNA-binding and transactivation domains in particular (Knight et al., 2005). The sequence of both *tfap2a* and *tfap2b* zebrafish genes are conserved with their respective vertebrate orthologues as well (Fig. S1) (Knight et al., 2005).

To further investigate the spatiotemporal expression domains of *tfap2a* and *tfap2b* within the developing renal field, we performed WISH with RNA antisense riboprobes over the time span of nephrogenesis. We found that *tfap2a* and *tfap2b* transcripts were expressed broadly in renal progenitors at the 10 somite stage (ss), however this expression became dynamically restricted to the distal region of the pronephros by the 28 ss (Fig. 2A). Through fluorescent *in situ* hybridization (FISH) studies, we confirmed that *tfap2a* and *tfap2b* were robustly co-expressed in nearly identical renal progenitor domains at the 10 ss (Fig. 2B). FISH at the 20 ss and 28 ss revealed that *tfap2a* had a mostly broader expression pattern across the distal pronephros compared to *tfap2b* (Fig. 2B), consistent with expression in the DE and DL segments. Differential *tfap2a/b* expression was noted rostrally and in the posterior pronephric duct region at the 20 ss and 28 ss, where only *tfap2a* transcripts were detected (Fig. 2B). Next, we sought to validate if *tfap2a* was expressed in the DE segment domain, which is demarcated by *slc12a1* expression, given the *trm* mutant phenotype (Wingert et al., 2007). *tfap2a* and *slc12a1* transcripts were co-localized at the 28 ss (Fig. 2C). These results indicate that *tfap2a* and *tfap2b* expression occurs in renal progenitors during a developmental window that positions them as possible participants in distal nephron development.

**Figure 2:**
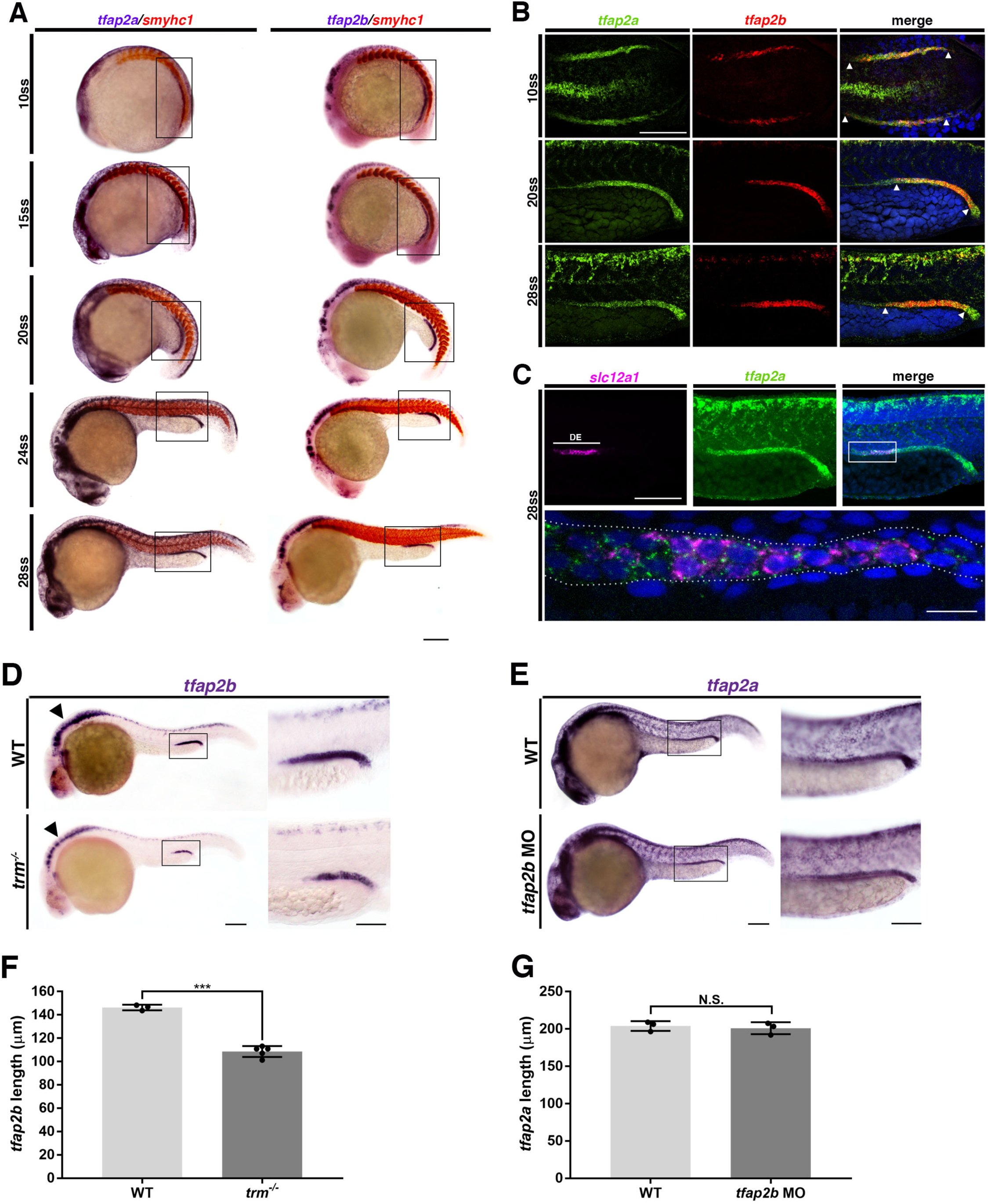
*tfap2a* and *tfap2b* transcripts are expressed in dynamic overlapping domains in developing nephrons, where *tfap2a* acts upstream of *tfap2b*. **A.** WISH of WT *tfap2a* and *tfap2b* expression (purple) at the 10, 15, 20, 24, and 28 ss. *smyhc1* (red) was used to mark somites. Black boxes indicate *tfap2* expression domains within developing renal progenitors. Scale bar = 200 µm. **B.** Double FISH of WT *tfap2a* (green) and *tfap2b* (red) transcript expression at 10 ss (flat mount), 20 ss, and 28 ss (lateral views). DAPI (blue) labels nuclei. White arrowheads demarcate cellular regions of overlapping transcripts within the pronephros. Scale bar = 70 µm. **C.** Double FISH of *slc12a1* (pink) and *tfap2a* (green) in WT embryo at 24 hpf. White box indicates area featured in bottom panel (60x z-stack). White dotted line outlines nephron tubule. DAPI (blue) labels nuclei. Scale bars = 70 µm, 10 µm. D. WISH analysis of *tfap2b* expression in WT and *trm*^-/-^. Arrowhead indicates differential hindbrain expression of *tfap2b*. Black box designates *tfap2b* expression within pronephros. Scale bars = 100 µm, 50 µm. **E.** WISH analysis of *tfap2a* expression in WT and *tfap2b* MO. Black box designates *tfap2a* expression within pronephros. Scale bars = 100 µm, 50 µm. **F.** Quantification of absolute length measurements of *tfap2b* expression domain within pronephros. **G.** Quantification of absolute length measurements of *tfap2a* expression domain within pronephros. n = 3 for each control and test group. Absolute measurements (in microns) were analyzed by unpaired t-tests. Data are represented as ± SD. ***p < 0.001, N.S. = not significant.

To explore potential genetic relationships between *tfap2a* and *tfap2b,* we performed loss of function experiments. *trm* exhibited significantly reduced *tfap2b* expression in the distal pronephros and the hindbrain region (Fig. 2D). Conversely, knockdown of *tfap2b* using a morpholino (MO) strategy revealed that *tfap2a* expression was unaffected throughout the embryo, including the pronephros (Fig. 2E). The *tfap2b* MO tool was verified to interrupt splicing by RT-PCR, which revealed that it caused inclusion of intronic sequence that encoded a premature stop codon, which is predicted to generate a truncated peptide (Fig. S3). In combination, these genetic studies suggest a more dominant role of *tfap2a* in the context of nephrogenesis, placing *tfap2a* upstream of *tfap2b*.

Because *tfap2a* and *tfap2b* have been demonstrated to function redundantly in the development of other tissue types, we next wanted to determine if these two factors could act similarly during nephrogenesis (Knight et al., 2005; Van Otterloo et al., 2018; Seberg et al., 2017; Bassett et al., 2012; Jin et al., 2015). To interrogate this, we performed combination knockdown studies and assayed a set of solute transporters that characterize the distal nephron segments at 24 hpf (Fig 3A). *tfap2b* deficiency alone had no detectable affects on distal solute transporter expression. *trm* and *tfap2a* morphants exhibited significant reductions in *slc12a1*, *slc12a3*, and *clcnk* expression as compared to WT (Fig. 3A, B, C, D). At 4 dpf, *trm* mutants still failed to express distal early solute transporters *slc12a1* and *romk2* (Fig. S4). However, the pronephros was functional between 2 and 3 dpf, based on assessment of renal clearance, which normally initiates during this time period, thereby ruling out developmental delay (Fig. S4). Knockdown of *tfap2b* in *trm* mutants caused a more severe *slc12a3* reduction than *tfap2a* deficient embryos (Fig. 3). Interestingly, there was not a statistically significant reduction in the *slc12a1* or *clcnk* domain length in *tfap2b* injected *trm* mutants versus *tfap2a* deficiency alone (Fig. 3). By comparison, *tfap2a/2b* morphants had statistically significant reduction of the *slc12a1*, *slc12a3*, and *clcnk* pronephros expression domains versus *tfap2a* deficiency alone. Notably, *tfap2a* morpholino targets all three splice variants, however the *trm* mutation only affects one of these splice variants (Fig. 1C). In light of this phenotypic spectrum, we concluded that the development of the distal nephron program is sensitive to the dosage of functional *tfap2a/tfap2b* alleles that are present. Taken together, these genetic studies reveal that the concerted action of *tfap2a* and *tfap2b* is necessary to fully turn on distal solute transporter programs, where *tfap2a* plays a more prominent role in this process upstream of *tfap2b*.

**Figure 3:**
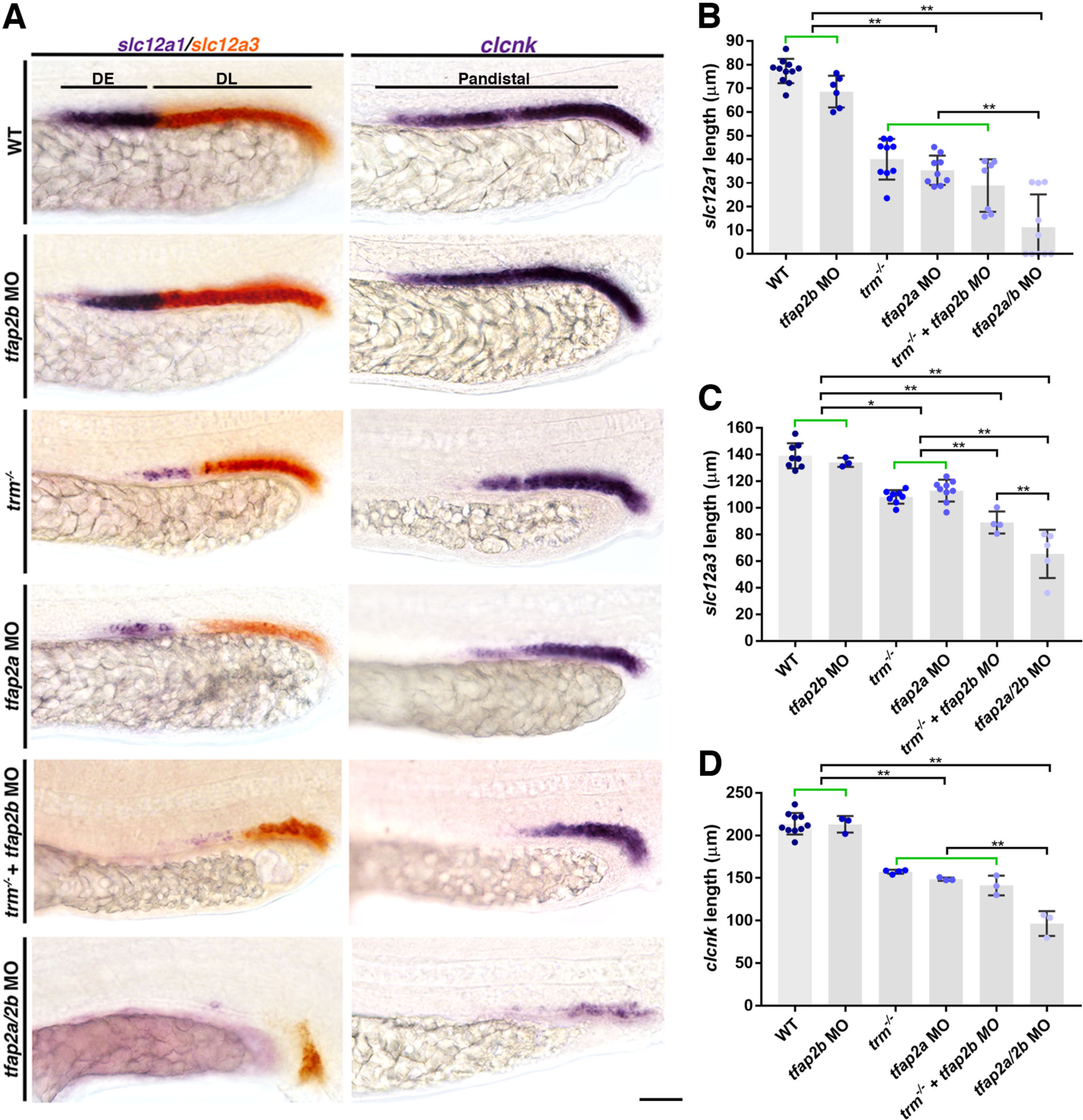
*tfap2a* and *tfap2b* function redundantly to activate distal nephron solute transporter signature. **A.** WT and *trm*^-/-^ embryos were microinjected with combinations of *tfap2a* and *tfap2b* splice-MOs. WISH was used to stain embryos for *slc12a1* (DE, purple), *slc12a3* (DL, red), and *clcnk* (Pandistal, purple) at 24 hpf. Black bars indicate WT marker domains. Scale bar = 35 µm. **B.** Quantification of absolute length of *slc12a1* expression domain. **C.** Quantification of absolute length measurements of *slc12a3* expression domain. **D.** Quantification of absolute length measurements of *clcnk* expression domain. n ≥ 4 for each control and test group. Measurements were compared by ANOVA. Data are represented as ± SD. *p < 0.05, **p < 0.01, Green brackets indicate not statistically significant.

### *tfap2a* is necessary and sufficient for DE differentiated cell expression signature

Next, we wanted to determine if provision of WT *tfap2a* transcripts could specifically rescue the absence of DE solute transporter expression in *trm*. Activation of a heat-shock inducible *tfap2a* transgene at the 8 ss restored *romk2* expression in *trm* mutants comparable to WT levels based on absolute length measurements (Fig. 4A). This result further underscores the conclusion that *tfap2a* deficiency is the single, specific cause of the *trm* phenotype. We then performed *tfap2a* gain of function studies by two independent methods: 1) employing an inducible *hs:tfap2a* transgenic line and 2) microinjection of *tfap2a* mRNA in WT embryos. When we overexpressed *tfap2a* by these approaches there was a significant expansion of the *romk2* expression domain which normally marks the DE segment (Fig. 4B). All of the heat-shock treated *hs:tfap2a* transgenic embryos exhibited an expanded *romk2* domain, while control non-heat-shocked embryos developed normal *romk2* domains. About 9% (12/132) of the *tfap2a* cRNA injected clutches presented with an increased *romk2* domain, however about 64% (85/132) of embryos were scored as dysmorphic. This lower phenotype penetrance, compared to the transgenic overexpression model, is likely caused by the toxic affects of *tfap2a* during early development, which has been reported to disrupt gastrulation (Li and Cornell, 2007). Interestingly, ectopic *romk2^+^* cells appeared to invade both the adjacent proximal and distal segment domains in these gain-of-function experiments.

**Figure 4:**
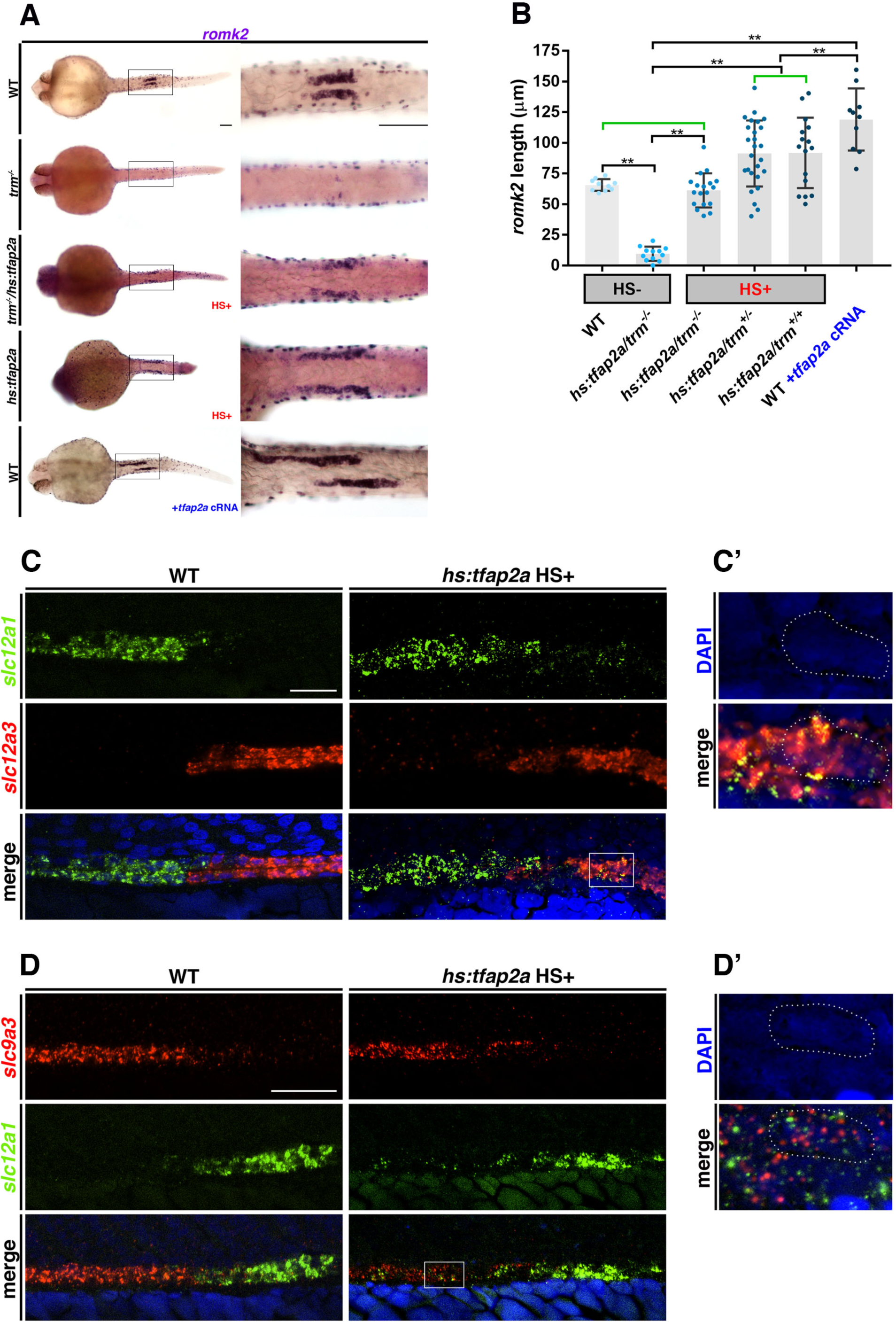
*tfap2a* is necessary and sufficient to drive the DE gene expression program. **A.** *trm*^-/-^*/hs:tfap2a* and *hs:tfap2a* were heat-shocked at the 8 ss to overexpress WT Tfap2a protein for rescue and gain of function studies. WT embryos were microinjected with *tfap2a* cRNA for an independent gain of function studies. Control and experimental samples subjected to WISH analysis for *romk2* (DE) expression at 24 hpf. Black bar indicates WT *romk2* domain. Scale bars = 70 µm. **B.** Quantification of absolute length measurements of *romk2* expression domain. n ≥ 4 for each control and test group. Measurements were compared by ANOVA. Data are represented as ± SD. **p < 0.01, Green brackets indicate not statistically significant. HS+ (red) signifies application of heat-shock. HS- (black) indicates no heat-shock. +*tfap2a* cRNA (blue) represents microinjection of *tfap2a* capped RNA at the 1-cell stage. **C.** Double FISH analysis of *slc12a1* (DE, green) and *slc12a3* (DL, red) in WT and heat shock-treated *hs:tfap2a* embryos at 24 hpf. Scale bar = 20 µm. White box indicates area imaged at higher (60X) objective in C’. **C’.** DAPI only (above) and merged channels (below). Dotted white line outlines a single cell coexpressing *slc12a1* and *slc12a3* transcripts. **D.** Double FISH analysis of *slc9a3* (panproximal, red) and *slc12a1* (DE, green) in WT and heat shock-treated *hs:tfap2a* embryos at 24 hpf. Scale bar = 35 µm. White box indicates area imaged at higher (60X) objective in D’. **D’.** DAPI (above) and merged channels (below). Dotted white line outlines a single cell coexpressing *slc9a3* and *slc12a1* transcripts. DAPI (blue) labels nuclei.

To examine more closely if *tfap2a* overexpression was inducing neighboring nephron segments to convert to a DE program, we performed double FISH on heat-shocked *hs:tfap2a* embryos. We detected *slc12a1^+^* cells within the *slc12a3*^+^ DL domain (Fig. 4C). Upon closer analysis, these cells were found to coexpress both *slc12a1* and *slc12a3* transcripts (Fig. 4C’). Heat-shocked *hs:tfap2a* animals also possessed *slc12a1^+^* cells spanning across the proximal domain that were coexpressing *slc9a3 (* Fig. 4D, D’). These phenotypes greatly contrast the WT situation, where there are sharp, clear boundaries between neighboring segment domains in the nephron (Fig. 4C, D). These results indicate that *tfap2a* overexpression is sufficient to sway the differentiation profile of proximal and distal late cell types by triggering the misexpression of DE-specific solute transporters.

### *tfap2a* drives DE terminal differentiation program

Previous studies have demonstrated *tfap2a* can regulate terminal differentiation of various cell types including migratory neural crest, melanocytes, statoacoustic ganglion neurons, noradrenergic neurons, and the trophoblast lineage (Barrallo-Gimeno et al., 2003; Kantarci et al., 2015; Seberg et al., 2017; Greco et al., 1995; Kim et al., 2001; Pfisterer et al., 2001; Handwerger, 2009). This literature, in light of our loss of function and gain of function results, led us to hypothesize that *tfap2a* controls the terminal differentiation of distal nephron cells. To explore this notion, we first wanted to determine if the nephron segments were patterned correctly in *trm* mutants.

To assess the pattern formation of nephron segments in *trm* mutants, we performed double WISH to assess the segment domains located adjacent to the DE, in this case the pan-proximal and DL. *trm* mutants exhibited a domain of *slc9a3* expression comparable to WT embryos (Fig. 5A). In both WT and *trm*, this *slc9a3^+^* region was followed by a gap situated at the position normally occupied by the DE segment, and the *slc12a3* expression domain, which is smaller in *trm* mutants, immediately followed this gap (Fig. 5A). The intact sequence of the pan-proximal, gap/placeholder, and then the DL segment suggested that the DE segment ‘footprint’ was present in *trm* mutants, and thus that pattern formation had proceeded during nephrogenesis (Fig. 5A). Additionally, *trm* mutants exhibited no alterations in *slc20a1a* or *trpm7* expression domains, which mark the PCT and PST segments, respectively (Fig. S5). *tfap2a* morphants also developed normal proximal segments, as well as the DE footprint (data not shown). These results indicate that *tfap2a* deficient embryos undergo normal segmental patterning of the nephron tubule.

**Figure 5:**
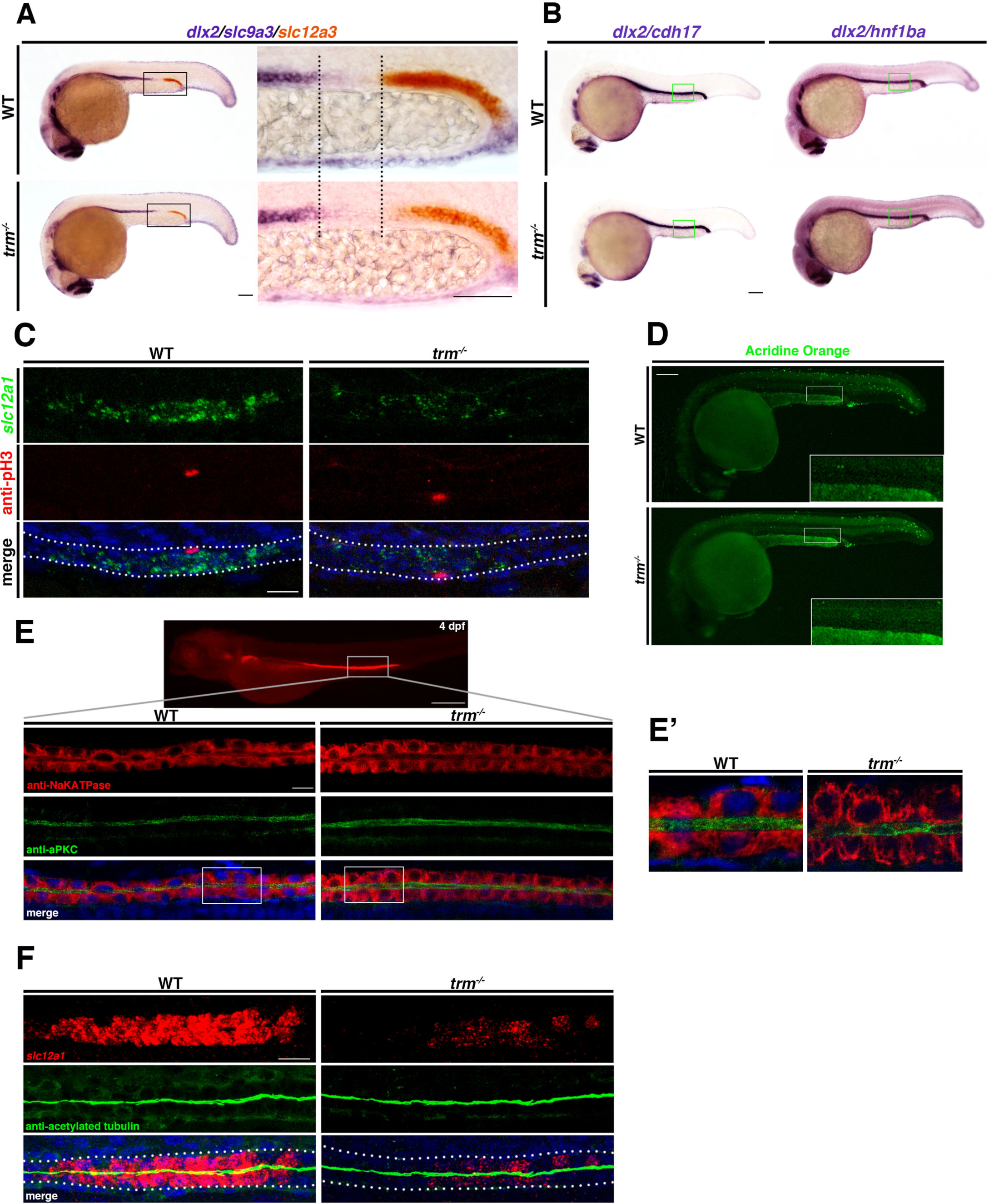
*tfap2a* is essential for the induction of terminal differentiation, but not cell proliferation, survival, polarity or ciliogenesis within the distal nephron. **A.** WISH used to stain *dlx2* (pharyngeal arches, purple), *slc9a3* (panproximal, purple), and *slc12a3* (DL, red) performed on 24 hpf WT and *trm*^-/-^ embryos to assess nephron patterning. Black dotted lines indicate presumptive area occupied by DE progenitors. Scale bars = 70µm. **B.** WISH analysis of *dlx2* and renal specification markers that span entire tubule (*cdh17* and *hnf1ba*) in WT and *trm*^-/-^ at 24 hpf. Green boxes indicate continuous expression of tubule markers in DE. Scale bar = 70 µm. **C.** Whole mount FISH and IF to visualize proliferating DE cells (*slc12a1*, green) in WT and *trm*^-/-^ at 24 hpf. anti-ph3 (red) labels proliferation. White dotted lines outline pronephric tubule (bottom). Scale bar = 10 µm. **D.** Acridine orange assay reveals no detectable difference in dying cell number (green) in WT and *trm*^-/-^ at 24 hpf. White box indicates inset (optical zoom) of distal nephron area. Scale bar = 70 µm. **E.** Survey of epithelial polarity proteins by whole mount IF in WT and *trm*^-/-^ at 4 dpf. anti-Na, K-ATPase (red) was used as a basolateral marker and anti-aPKC (green) was used as an apical marker. Top image represents WT Na, K-ATPase protein expression in 4 dpf. White boxes indicate region in E’. Scale bars = 200 µm, 10 µm. **E’.** Regions highlighted in B showing normal protein localization. **F.** Whole mount FISH with IF to assess cilia (anti-acetylated tubulin, green) morphology in the DE (*slc12a1,* red) of WT and *trm*^-/-^ at 24 hpf. White dotted lines demarcate nephron. Scale bar = 10 µm. DAPI (blue) labels nuclei.

We then examined development of the corpuscle of Stannius in *trm* mutants, which is an endocrine gland situated between the DE and DL (Cheng and Wingert, 2015). We used WISH to assess the expression of transcripts encoding *stanniocalcin 1* (*stc1*), a specific CS marker. Compared to WT embryos, *trm* mutants exhibited severely reduced *stc1* expression (Fig. S5). In sum, these data rule out the occurrence of possible fate switches with adjacent nephron cell types that could account for the loss of the DE marker expression, and suggest that in addition to the DE, *tfap2a* may regulate CS lineage development and/or CS differentiation.

Next, we wanted to determine if *trm* mutant cells occupying the DE region were specified as kidney. To do this, we individually assayed for the expression of genes that are expressed robustly throughout the entire nephron tubule. Identical to WT embryos, *trm* mutants showed no gaps in expression of *cdh17* and *hnf1ba*, illustrating that mutant DE cells were fated to a kidney lineage identity (Fig. 5B). Further, we assessed if any alterations in cell proliferation or cell death occurs in response to loss of *tfap2a*. We detected no visible changes in the number of pH3^+^ cells within the DE domain of *trm* mutants compared to WT controls at 24 hpf (Fig. 5C). Additionally, there were no perceivable differences in the number of dying cells labeled with acridine orange in the distal pronephros compared to WTs at 24 hpf (Fig. 5D).

Renal progenitor differentiation in the zebrafish pronephros is known to entail an MET of the mesenchymal progenitors, with establishment of polarity and lumen formation, as well as changes in cellular organelles such as cilia. Therefore, we sought to determine whether *trm* mutant DE cells exhibited any of these various differentiated features of the nephron tubular epithelium. Differentiated pronephros cells exhibit apical-basal polarity and form either a primary cilium or multiple cilia by the 24 hpf stage (Gerlach and Wingert, 2013, 2014; McKee et al., 2014; Marra and Wingert 2016; Marra et al., 2016). We performed whole mount IF to analyze the expression of basolateral marker Na, K-ATPase and the apical adaptor complex aPKC in the distal early region at 4 dpf. We found Na, K-ATPase and aPKC proteins were properly localized in *trm* mutants as compared to WT, therefore indicating that epithelial polarity was correctly established within the nephron tubules (Fig. 5E, E’). Further, *trm* mutants had a clearly discernible tubule lumen, indicating that tubulogenesis had proceeded analogous to WT embryos (Fig. 5E, E’). Next, we combined FISH of *slc12a1* with whole mount IF of acetylated tubulin to determine if cilia formation occurs within the mutant DE segment region. At 24 hpf, cilia arrangement and morphology in *trm* mutants was comparable to WT (Fig. 5F). This indicates that cilia assembly occurred normally in the mutant DE cells, which were visualized based on their nearly abrogated *slc12a1* signal (Fig. 5F). Taken together, these data indicate that mutant DE cells exhibit mature epithelial qualities, however cannot fully turn on specific solute transporters, which are indicators of terminal differentiation and ultimately dictate segment-specific physiological functions. From this, we conclude that *trm* mutants exhibit a unique block in the terminal differentiation of distal nephron cells, which involves the acquisition of segment-specific solute transporter proteins but is not linked to MET, polarity establishment, tubulogenesis or ciliogenesis programs.

### *tfap2a* functions downstream of *irx3b* and upstream of *irx1a* in the distal pronephros

We next wanted to understand the genetic relationship of *tfap2a* with known segment patterning factors, to establish the hierarchical regulation of *tfap2a* during distal segment development. Previous studies have shown *Irx3/irx3b* are required for the development of *slc12a1*^+^ distal tubule cells in *Xenopus* and zebrafish, respectively (Wingert and Davidson, 2008; Reggiani et al., 2007; Marra and Wingert, 2014). Because of this requirement, we first selected *irx3b* as a putative *tfap2a* gene regulatory network candidate for investigation.

To determine if *tfap2a* and *irx3b* are co-expressed during pronephros development, we performed double FISH studies. We found *tfap2a* and *irx3b* transcripts were co-localized in developing distal nephron cells at the 20 ss (Fig. 6A). Because of the *tfap2a/irx3b* overlapping expression patterns, we rationalized that these factors could be interacting in the same developmental pathway. Therefore, we performed knockdown experiments to determine potential pathway interactions between *tfap2a* and *irx3b*. Upon *tfap2a* knockdown, the *irx3b* expression domain was unchanged (Fig. 6B). However, upon *irx3b* knockdown, the *tfap2a* expression domain was significantly truncated (Fig. 6B, C). The regional loss of *tfap2a* transcripts in *irx3b* morphants equates to the DE segment address. We also observed a loss of *tfap2a* expression in migrating neural crest streams in *irx3b* morphants (Fig. 6B). To further validate if *tfap2a* acts downstream of *irx3b*, we performed rescue experiments in *irx3b* morphants. Overexpression of the *hs:tfap2a* transgene was unable to rescue *romk2* expression in *irx3b* knockdowns (data not shown). We postulate this is because *irx3b-* deficienc*y* causes loss of *hnf1ba* expression within the DE progenitor compartment, therefore these cells are not competent to respond to Tfap2a activity (Naylor et al., 2013). These results suggest that *tfap2a* activates the DE program downstream of *irx3b*.

**Figure 6:**
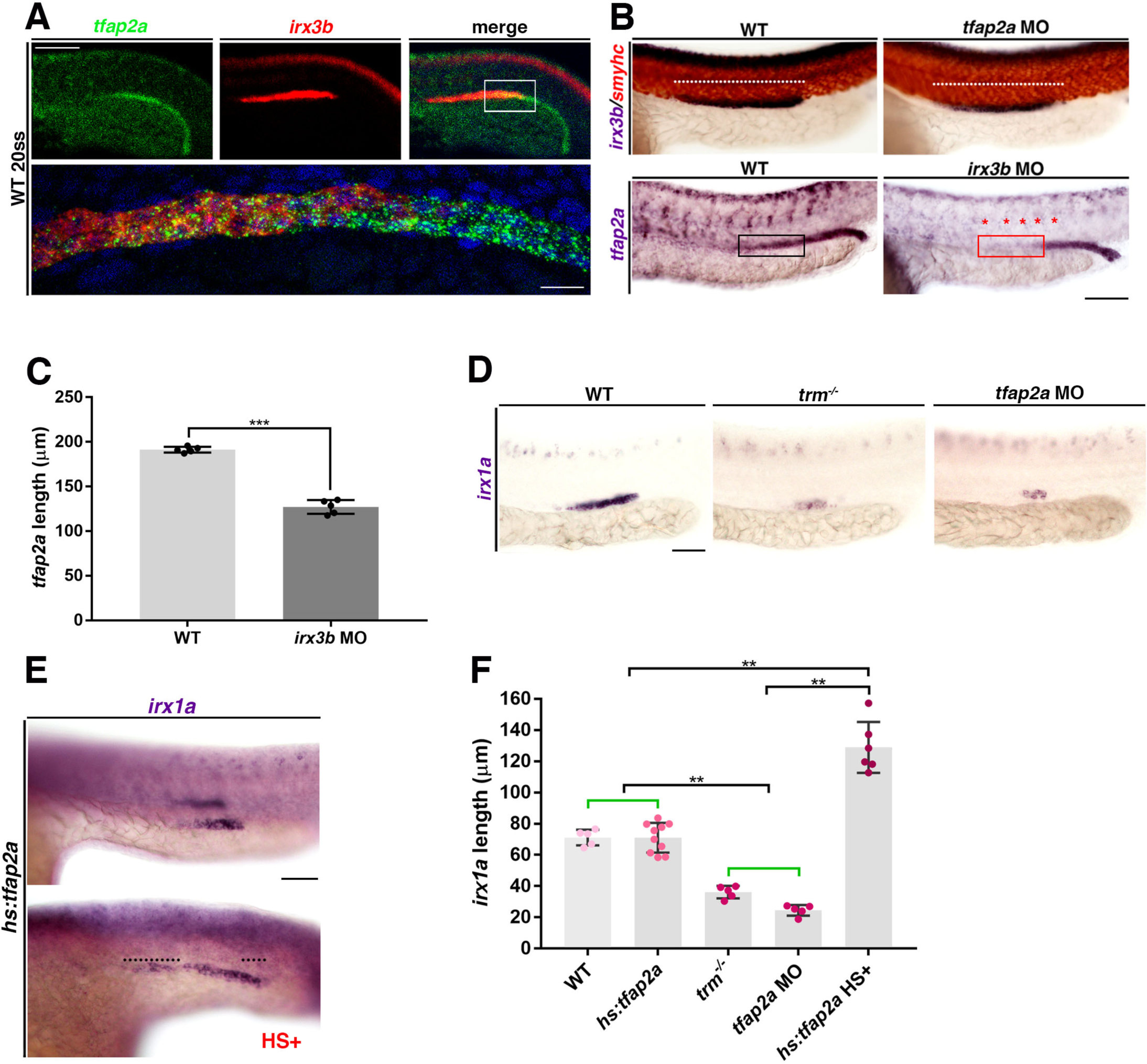
*tfap2a* interplays with the Iroquois homeobox genes *irx3b* and *irx1a* during nephrogenesis. **A.** Whole mount double FISH in 20 ss WT animals reveal *tfap2a* transcripts (green) and *irx3b* transcripts (red) are coexpressed distal pronephros. White box focuses on area of intense coexpression. Bottom panel indicates area outlined by white box at 60X magnification. DAPI (blue) labels nuclei. Scale bars = 70 µm, 10 µm. **B.** Top panel: WISH for *irx3b* (purple) and *smyhc1* (red) in WT and *tfap2a* morphants at 24 hpf. White dotted line indicates *irx3b* expression domain, which does not change due to *tfap2a*-deficiency. Bottom panel: WISH for *tfap2a* expression in *irx3b* morphants at 24 hpf. Black box indicates presence of *tfap2a* transcripts in DE segment domain in WT. Red box indicates absence of *tfap2a* transcripts in DE segment domain in *irx3b* morphants. Red asterisks (*) indicate loss of *tfap2a* expression within the neural crest streams in *irx3b* morphants. Scale bar = 70µm. **C.** Quantification of *tfap2a* expression domain length. Measurements were compared by unpaired t-test. Data are represented as ± SD. ***p<0.001. **D.** WISH reveals reduced *irx1a* expression in *trm*^-/-^ and *tfap2a* morphants as compared to WT at 24 hpf. Scale bar = 35 µm. **E.** WISH of *irx1a* expression in *hs:tfap2a* untreated and heat shock-treated (red HS+) at 24 hpf. Black dotted lines denotes increased expression of *irx1a* in heat shock-treated *hs:tfap2a* embryos. Scale bar = 35µm. **F.** Quantification of *irx1a* expression domain length in WT, *hs:tfap2a* (untreated), *trm*^-/-^, *tfap2a* morphants, and heat shock treated (red HS+) *hs:tfap2a*. Measurements were compared by ANOVA. Data are represented as ± SD. **p < 0.01, Green brackets indicate not statistically significant.

Previous literature has determined *Irx1* and *Irx3* are dually required for *Xenopus* pronephros development (Reggiani et al., 2007; Alarcon et al., 2008). Importantly, loss of *Irx1* in *Xenopus* results in proximal downregulation of the *Nkcc2* expression domain, the *slc12a1* equivalent in zebrafish (Reggiani et al., 2007). Therefore, we chose Iroquois homeobox family member, *irx1a*, as another important molecular player for analysis. In WT embryos, *irx1a* transcript expression was primarily localized to the DE segment (Fig. 6D) (Cheng et al., 2001). In *trm* mutants and *tfap2a* morphants, *irx1a* expression was nearly abrogated, with only a few remaining nephron cells expressing transcripts (Fig. 6D). When we induced overexpression of *tfap2a* at the 8 ss, the *irx1a* expression domain length was significantly expanded, indicating that *tfap2a* functions to activate *irx1a* expression directly or indirectly (Fig. 6E, F). Our results place *tfap2a* function with respect to *irx* activity, indicating that *tfap2a* functions downstream of *irx3b* and upstream of *irx1a* during distal segment differentiation. Taken together, our genetic analyses suggest a working model in which *tfap2a* coordinates a genetic regulatory network, likely through direct and indirect interactions, that controls the differentiation of distal nephron progenitors (Fig. 7).

**Figure 7:**
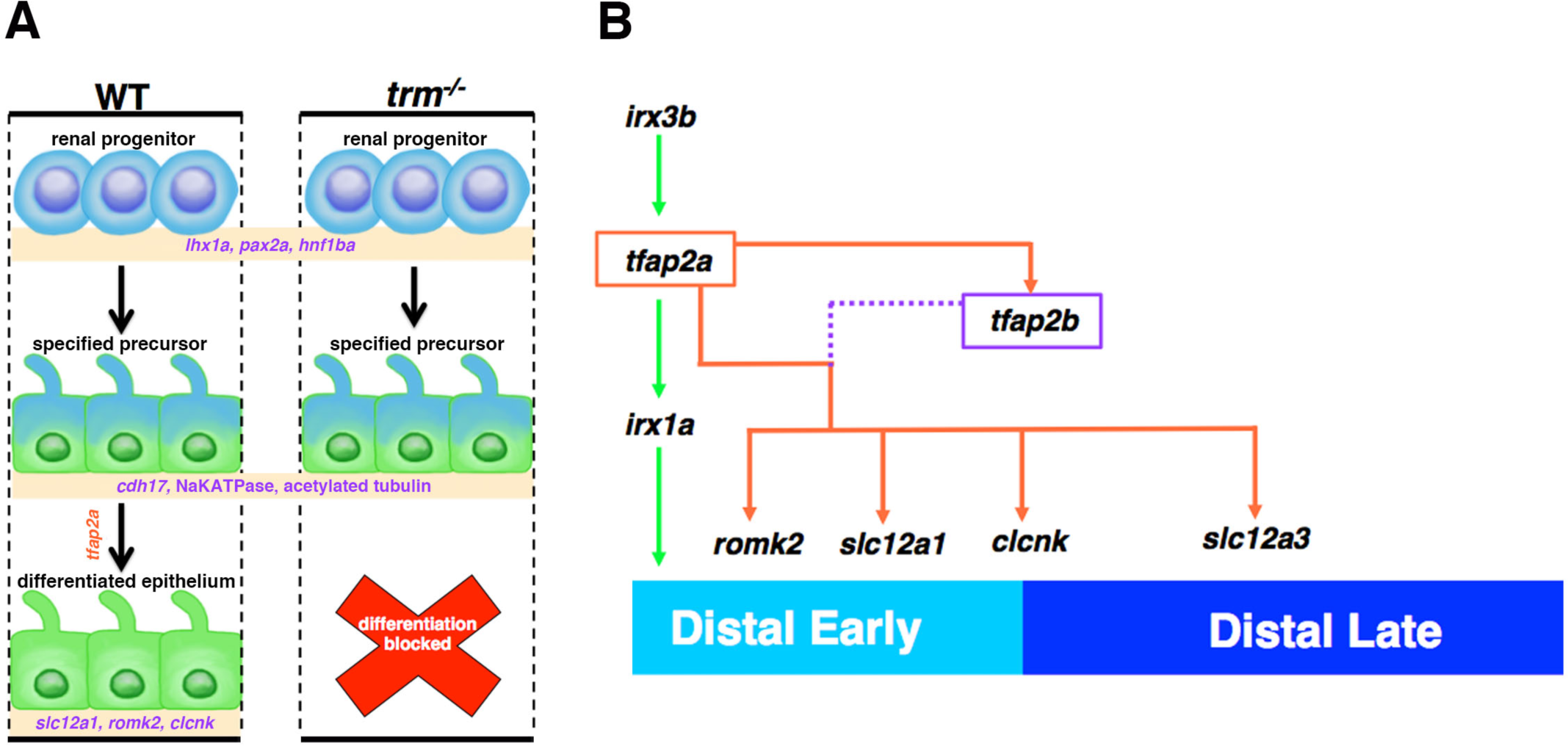
*tfap2a* and *tfap2b* function in a proposed genetic regulatory network to control distal nephron differentiation. **A.** Schematic comparing DE progenitor maturation in WT and *trm*^-/-^. Mutant cells display no perturbations in the early specification of the renal lineage (marked by *lhx1a*, *pax2a*, and *hnf1ba*). Mutant progenitors undergo segment specification and exhibit features of mature epithelium (*cdh17*, Na, K-ATPase, and acetylated tubulin). In the final phase of differentiation, mutant cells fail to express DE-specific solute transporters (*slc12a1*, *romk2*, and *clcnk*). B. Diagram depicts *tfap2a* distal nephron GRN. *irx3b* promotes *tfap2a* expression, and *tfap2a* functions upstream of *irx1a* (green arrows). *tfap2a* acts upstream of *tfap2b* as the core regulator of solute transporter expression (orange arrows). *tfap2b* functions redundantly (purple dotted line) to activate distal solute transporters (*romk2*, *slc12a1*, *clcnk*, and *slc12a3*).

## Discussion

Here, we have shown that *tfap2a* is required for distal nephron segment differentiation. We propose a model in which *trm* mutant cells progress through normal nephron developmental checkpoints until the final stage of differentiation (Fig. 7A). Our data supports the conclusion that *tfap2a* deficiency does not affect the derivation of the kidney lineage from intermediate mesoderm, as the expression of *hnf1ba* and *cdh17* are unaffected. Further, we found that *trm* undergo nephron specification and epithelialization, as *tfap2a*-deficient cells exhibit segmental patterning, proper localization of polarity proteins, and form cilia. However, mutant cells appear to be frozen nonetheless in a specified renal precursor state, as they fail to turn on the suite of terminal differentiation genes that encode the distal solute transporters *slc12a1*, *romk2*, and *clcnk*. Taken together, we conclude that *tfap2a* is required for a discrete genetic circuit during terminal differentiation of the distal nephron epithelium.

Interestingly, this discovery disentangles the control of the solute transporter transcriptome from other differentiation processes, such as MET and polarity establishment in renal progenitors. Additionally, we have assembled a proposed Tfap2 genetic circuit that functions to achieve differentiation of distal nephron epithelium within the zebrafish embryonic kidney (Fig. 7B). In this network, *tfap2a* functions upstream of *tfap2b* as indicated by our genetic analyses. However, both *tfap2a* and *tfap2b* function synergistically in renal progenitors to turn on distal solute transporter genes, a level of redundancy that likely serves to amplify and reinforce this specific differentiation signal. This model is supported by the findings that elimination of *tfap2a* alleles led to defects in solute transporter expression, however elimination of *tfap2b* alleles alone had no consequence. Compound knockdown of *tfap2a* and *tfap2b* yielded the most severe phenotype, unveiling this layer of functional redundancy. As the *trm* mutation affects only one of the *tfap2a* spliceoforms, it is interesting that disrupting only this transcript results in a kidney phenotype, and suggests it encodes the sole protein variant that is fundamental for kidney development. The current study does not reconcile whether *tfap2a* and *tfap2b* are interacting by direct or indirect modes of regulation, and whether the suite of targets are direct or indirect (Fig. 7B). For example, it is possible *tfap2a* binds to the *tfap2b* promoter region functioning as a transcriptional activator. It is also feasible Tfap2a heterodimerizes with Tfap2b affecting the transcription of downstream target genes accordingly. These potential biochemical mechanisms are crucial areas for future investigation.

Further, our genetic model supports the conclusion that *tfap2a* acts in the same pathway as Iroquois homeobox genes *irx3b* and *irx1a*. Our genetic experiments indicate *irx3b* promotes *tfap2a* expression, and *tfap2a* functions upstream of *irx1a* (Fig. 7). These genes have been previously implicated as necessary for intermediate-distal early nephron development in zebrafish and frogs (Wingert and Davidson, 2011; Reggiani et al., 2007). Importantly, Iroquois factors likely play conserved roles in mammalian counterparts, as *Irx3* and *Irx1* define intermediate segment territories in developing S-shaped bodies (Reggiani et al., 2007). Defining whether Irx3b directly regulates *tfap2a*, and if Tfap2a directly regulates *irx1a*, will also be important to discern in future studies as well.

Additionally, we discovered an intriguing phenotype when we globally overexpressed *tfap2a*, where *slc9a3*^+^ proximal tubule and *slc12a3*^+^ distal late tubule cells ectopically coexpress *slc12a1*, a marker of the DE tubule program (Fig. 5). This indicates that renal progenitors are competent to respond to Tfap2a, which is sufficient to activate the DE differentiation circuit. These mixed segment identities induced by *tfap2a* gain of function parallels a phenotype recently described as ‘lineage infidelity,’ which was observed in differentiating nephrons of Hox9,10,11-knockout mice (Magella et al., 2018; Drake et al., 2018). These studies similarly found individual cells that were dually expressing markers of more than one nephron segment. Specifically, in E18.5 *Hoxa9,10,11/Hoxd9,10,11*-deficient kidneys, cells were scattered throughout Hnf4a^+^ proximal tubules co-expressing collecting duct markers DBA and Krt8. Strikingly, mutant kidneys sometimes possessed entire Slc12a1^+^ ascending loop of Henle (the DE analog in zebrafish) domains that co-labeled with Krt8. These developing mutant nephrons undergo a normal segment specification phase (E15.5), but later fail to maintain appropriate differentiation programs. It is an interesting prospect that Hox mutants exhibit a normal specification phase, as our data similarly suggests that *tfap2a* is dispensable for nephron patterning, but necessary for inducing a proper differentiation state. During zebrafish neural crest development, *tfap2a* promotes expression of Hox group 2 genes to form segments of the pharyngeal skeleton (Knight et al., 2003; Barrallo-Gimeno et al., 2003). Collectively, these studies warrant Hox genes as important future areas of study regarding the Tfap2 genetic regulatory network controlling nephron differentiation.

With the advent of next-generation sequencing technologies, recent work in the field has identified new targets within the *Tfap2* genetic regulatory network (Seberg et al., 2017). To do this, microarray analyses and ChIP-seq were employed in tandem to identify novel players regulating melanocyte development within the *Tfap2a* GRN. To collect candidates, microarray analysis of *tfap2a*^-/-^ zebrafish trunks and *tfap2a-* deficient mouse melanocyte lines were conducted. Microarray results were compared to ChIP-Seq data gathered from mouse and human melanocytes to determine which differentially regulated genes were direct transcriptional targets of *Tfap2a*. The findings suggested *Tfap2a* directly regulates effectors of melanocyte terminal differentiation (e.g. *dct, mlpha, mc2r, sox10, mitf*). Conducting microarray analysis of our *trm* mutant zebrafish embryos and overlapping this data with the previously published mammalian kidney Tfap2a Chip-Seq data set (Pihlajamaa et al., 2014) will help to identify putative direct targets governing terminal differentiation of distal nephron cells.

A separate study performed RNA-sequencing analysis on dissected mandibular processes from double conditional *Tfap2a/Tfap2b* mouse mutant embryos to find *Tfap2a/2b* target genes during branchial arch patterning (Van Otterloo et al., 2018). Upon evaluating differentially expressed genes by Geneset enrichment analyses (GSEAs) ‘homeobox transcription factors’ was identified as the number one over-represented cluster. Further genetic workup of Dlx, Msx, and Emx homeobox gene families established these factors as major network targets under the control of Tfap2a/b during branchial arch development. Performing RNA-profiling and GSEAs of *tfap2a*-deficient developing nephrons would aid in pinpointing molecular pathways for future workup. The majority of differentially expressed genes in *Tfap2a/2b* murine branchial arches were associated with regions corresponding to poised histone marks, supporting a direct mode of regulation (Van Otterloo et al., 2018). However, it remains a possibility that *tfap2a* regulates transcription in nephron precusors via an intermediate factor or chromatin modifier as part of the GRN. In further support of an indirect mode of regulation, Tfap2a acts as a tissue-specific pioneer factor in the epididymis to modify chromatin structure activating androgen receptor signaling (Pihlajamaa et al., 2014). We speculate that Tfap2a/b operates both directly and indirectly to regulate expression of GRN components responsible for terminal differentiation of distal nephron tubules. To entertain this prospect, ATAC-seq and ChIP-seq methods could be installed in the future to determine differential chromatin accessibility in *tfap2a*-deficient renal progenitors.

*Tfap2a* has roles in induction, early specification, patterning, cell survival, and differentiation depending on the tissue type (Eckert et al., 2005). In neural crest, *tfap2a* plays dual roles in early development and later differentiation events. *tfap2a* and *foxd3* establish proper Bmp and Wnt signaling, which is required for early neural crest induction (de Croze et al., 2011; Bhat et al., 2012). In *tfap2a*-deficiency, migratory neural crest cells undergo increased apoptosis, indicating *tfap2a* is necessary for survival. It is speculated that because these cells cannot differentiate properly, they undergo apoptosis (Knight et al., 2003). Further, *tfap2a* is required for the specification and differentiation of inner ear neurons (Kantarci et al., 2015). *tfap2a* is also a necessary element for establishing preplacodal ectoderm competence and specification of ectoderm lineages (Bhat et al., 2012; Kwon et al., 2010). The concerted action of *tfap2a* and *phox2a* promote the differentiation of noradrenergic neurons in the central nervous system (Holzschuh et al., 2003). *tfap2a* also stimulates pathways that promote melanocyte terminal differentiation and survival (Van Otterloo et al., 2010; Seberg et al., 2017). *tfap2a* and *tfap2b* are required for the survival of sympathetic progenitors and differentiated sympathetic neurons (Schmidt et al., 2011). Differentiation of amacrine cells during retinogenesis is also induced by *tfap2a* and *tfap2b* (Bassett et al., 2012; Jin et al., 2015), and they function redundantly and non-autonomously to regulate cartilage patterning by modulating Fgf signals in the pharyngeal ectoderm (Knight et al., 2005). Interestingly, in the developing mouse kidney, *Tfap2b* is required for the maintenance and survival of renal epithelium (Moser et al., 1997). In *Tfap2b* null mice, distal tubules and collecting duct cells undergo a massive wave of apoptosis. Additionally, histology revealed mutant kidneys possess numerous cysts in the distal tubules and collecting ducts. Anti-apoptotic genes *bcl-X_L_*, *bcl-w*, and *bcl2* are strongly downregulated, supporting the idea that *Tfap2b* programs cell survival during embryogenesis. However, in our *trm* mutant, we are confident cell death does not contribute to the decreased solute transporter expression observed. Upon acridine orange analysis, we saw no obvious increase in the number of dying cells. Further, we observe no gaps in the nephron tubule as a result of dying cells, as indicated by continuous expression of *cdh17* expression at 24 hpf or at later time points. Our data strongly suggests that the main function of *tfap2a* and *tfap2b* during nephron development is not specification, patterning, or survival, but rather to promote terminal differentiation of distal nephron epithelium.

Based on recent studies in the zebrafish model, upstream candidates for regulating *tfap2a* and *tfap2b* may occupy the prostaglandin signaling pathway, which is essential to control the balance of DE and DL territories during zebrafish nephrogenesis (Poureetezadi et al., 2016), or include transcription factors like *mecom*, *tbx2a/2b, or emx1* that regulate DL development (Li et al., 2014; Drummond et al., 2016; Morales et al., 2018). Additional network candidates which may crosstalk with *tfap2a/2b* include *sall1* and *sox11*. In mammals *Sall1* has been found to be a critical factor in the development of the thick ascending limb (TAL), which is the segment analogous to the zebrafish DE region (Basta et al., 2017). *Sall1* mutants exhibited significantly decreased expression of *Kcnj1* (*Romk2*), *Slc12a1*, *Irx2*, and *Pou3f3*, among other major loop of Henle and distal lineage genes. Immunohistochemistry analysis revealed a near total loss of *Slc12a1*^+^ loop of Henle structures in the inner medulla of mutant embryos. In the murine kidney, *Sox11* is also necessary for proper loop of Henle ontogeny (Neirijnck et al., 2018). *Sox11*-deficient kidneys have significantly reduced expression of *Slc12a1*, *Irx1*, and *Irx2. Sall1* and *Sox11* are excellent candidates to situate in the Tfap2a GRN, due to their involvement in the DE/TAL segment development. Additionally, *Sall1*, *Sox11*, and *tfap2a* mutations all reduce Irx gene expression, suggesting they may act in the same pathway.

Knowledge about the terminal differentiation programs of each nephron segment has central importance for understanding kidney disease and to advance regenerative medicine. Human BOFS is associated with the occurrence of dysplastic kidneys, but the underlying mechanisms are not known. Our zebrafish *trm* mutant provides an opportunity to model aspects of BOFS at the molecular level of the nephron. With regard to kidney engineering, current groups face major challenges of generating mature, differentiated nephron structures in kidney organoid cultures (Hariharan et al., 2015; Chambers et al., 2016; Oxburgh et al., 2017; Takasato et al., 2017). However, growing mouse and particularly human kidney organoids is an immensely promising technology to study kidney development, model renal disease, and perform nephrotoxicity assays (Morizane and Bonventre, 2017). Reconstructing the mammalian nephron requires understanding the correct signals to guide stem cells down the appropriate differentiation paths to generate highly specialized compartments of cells. While terminally differentiated nephrons have yet to be achieved in organoid cultures, the discovery of terminal differentiation factors, like *tfap2a* and *tfap2b*, can herald progress in this crucial aspect of the kidney organoid field. In sum, our work indicates that further elucidation of the *Tfap2a/TFAP2A* gene regulatory network in zebrafish, murine, and human nephron progenitors can shed valuable insights into nephron differentiation and congenital renal disease.

## Materials and Methods

### Ethics statement and zebrafish husbandry

Adult zebrafish were maintained in the Center for Zebrafish Research at the University of Notre Dame Freimann Life Science Center. All studies were performed and supervised with by the University of Notre Dame Institutional Animal Care and Use Committee (IACUC), under protocol numbers 13-021 and 16-025. Tübingen strain animals were used for WT experiments. Zebrafish embryos were raised in E3 embryo media, staged, and fixed as described (Kimmel et al., 1995).

### Whole mount and fluorescent *in situ* hybridization (WISH, FISH)

WISH and FISH were performed as described (Marra et al., 2017; Brend and Holley, 2009; Lengerke et al., 2011; Cheng et al., 2014) with antisense RNA probes. Probes were synthesized using IMAGE clone template plasmids for *in vitro* transcription (Wingert et al., 2007; Wingert and Davidson, 2011). Digoxigenin-labeled probes consist of: *wt1b*, *slc20a1a*, *slc12a1*, *dlx2a*, *romk2*, *tfap2a*, *tfap2b*, *clcnk*, *slc12a3*, *slc9a3*, *cdh17*, *hnf1ba*, *irx3b*, *irx1a*, *trpm7*, and *stc1*. Fluorescein-labeled probes consist of: *tfap2b*, *slc12a1*, *slc12a3*, *tfap2a*. For all gene expression studies, each analysis was performed in triplicate with sample size of at least n=20 for each replicate. Representative animals were imaged and absolute length measurements were collected.

### Whole mount immunofluorescence (IF)

Whole mount IF studies were completed as described (McCampbell et al., 2015). To assess Tfap2a protein expression, anti-tfap2a (1:50) (LifeSpan Biosciences) and anti-goat secondary antibody were used. To analyze proliferation, anti-phospho-Histone H3 (1:200) (Millipore), and anti-rabbit secondary antibody (Alexa Fluor, Invitrogen) were used (Kroeger et al., 2017). For cilia studies, anti-acetylated α-tubulin (1:400) and anti-mouse secondary antibody (Alexa Fluor, Invitrogen) were used (Marra et al., 2017). Monoclonal α6F anti-NaKATPase (1:35) (Developmental Studies Hybridoma Bank) and anti-aPKC (1:250) (Santa-Cruz) were applied to embryos incubated in 0.003% PTU to prevent pigmentation and fixed in Dent’s solution (80% methanol, 20% DMSO) overnight at 4°C (Gerlach and Wingert, 2014). Anti-mouse and anti-rabbit secondary antibodies (Alexa Fluor, Invitrogen) were used respectively. All fluorescently-conjugated secondary antibodies were applied at a 1:500 dilution, and 4,6-diamidino-2-phenylindole dihydrochloride (DAPI) (Invitrogen) was used to stain nuclei.

### Image acquisition and statistical analysis

A Nikon Eclipse Ni with DS-Fi2 camera was used to image WISH samples. A Nikon C2 confocal microscope was used to image whole mount FISH and IF samples. The polyline tool in Nikon Elements imaging software was used to measure gene expression domains. A minimum of 3 representative samples for each control and experimental group were imaged and measured. Averages and standard errors were calculated. Unpaired t-tests or one-way ANOVA tests were completed for statistical analyses.

### Mutagenesis, whole genome sequencing and genotyping

WT zebrafish were exposed to ethylnitrosurea and haploids generated as described (Kroeger et al., 2014). Whole genome sequencing was performed as described (Leshchiner et al., 2012). Pools of 20 *trm* mutants and 20 WT siblings were identified by WISH analysis for *slc12a1* (DE) expression. DNA isolation was conducted using the DNAeasy blood and tissue kit (Qiagen). WGS results were interpreted by SNPtrack software (Leshchiner et al., 2012; Ryan et al., 2013). Isolation of genomic DNA from individual *trm* animals was performed and PCR amplification of the *tfap2a* locus was completed using the following primers: forward 5’-TTTGAACGCTGGCCACCGCCACCTCGCCCTACAATTATTGTTGGCTTGATTTAATTTGCACGTTCGTTTTTGATTTGTCCTTCTGAATTTCACGTCTTTT-3’ reverse 5’ AAATGTTTGGTTTTCGTTTACCAGTTAAAATCCTACCGAAAGGCAAAGGAAATTAACAATTAACCACAGCTCACATGAAGAAAATCTTTGTAATAGCCTT-3’. For all studies, *trm* mutants were confirmed by genotyping and/or abrogated *dlx2* expression. The QIAquick PCR Purification Kit was used to purify PCR product and sequenced with the forward primer by the University of Notre Dame Genomics Core Facility. Genotyping of *hs:tfap2a* transgenic (*Tg* (*hsp70:tfap2a*)^x24^, which was a generous gift from Bruce Riley, was conducted by performing PCR amplification (34 cyles, 60°C annealing) of the transgene and running product on a 1% agarose gel. The following primers were used: forward 5’-CTCCTCTCAATGACAGCTG-3’ reverse 5’-ATGGCGGTTGGAAGTCTGAA-3’.

### Overexpression experiments

To activate the heat-shock inducible *tfap2a* transgene, heterozygous transgenic embryos were incubated at 38°C for 30 minutes as described (Bhat et al., 2012; Kantarci et al., 2015). For rescue and gain-of-function studies, transgenic embryos were heat shocked at the 8ss. For FISH gain-of-function studies, transgenic embryos were heat shocked at the 10ss. For cRNA synthesis, the open reading frame of *tfap2a* was subcloned into the pCS2 vector. The primers used for subcloning were: forward 5’-GATCATCGATGCCGCCACCATGTTAGTGCACAGTTTTTCCGCGATGGATC-3’ reverse 5’-GATCTCTAGATCACTTTCTGTGCTTCTCATCTTTGTCACC-3’. For *in vitro* transcription, *tfap2a* template was linearized using Not1 restriction enzyme. Runoff reactions were performed using the mMessage mMachine Sp6 kit (Ambion). 50 picograms (pg) of *tfap2a* cRNA were injected into 1 cell stage embryos.

### Morpholino knockdown and RT-PCR

All morpholino oligonucleotides were synthesized by Gene Tools, LLC. *tfap2a* MO-splice (*tfap2a* MO4) targets the exon 2 – intron 2 splice site: 5’-AGCTTTTCTTCTTACCTGAACATCT-3’ [36]. *tfap2b* MO-splice (*tfap2b* MO1) targets the exon 4 – intron 4 splice site: 5’-GCCATTTTTCGACTTCGCTCTGATC-3’ (Knight et al., 2005). *irx3b* MO-ATG (*irx3b* MO2) targets the start site: 5’-ATAGCCTAGCTGCGGGAGAGACATG-3’. Morpholinos were solubilized in DNase/RNase free water to create 4mM stock solutions and stored at - 20°C. The stocks were diluted as follows for microinjection: *tfap2a*-MO 1:12, *tfap2b*-MO 1:10, *irx3b* MO 1:10. 1-cell stage embyros were injected with approximately 3 nl of morpholino. All splice-blocking MOs were verified by RT-PCR. Transcript analysis of *tfap2a* and *tfap2b* splicing in WT, WT sibs, *trm* mutants, *tfap2a* morphants, and *tfap2b* morphants was performed using RT-PCR (Galloway et al., 2008). In brief, RNA was isolated from pools of about 20 embryos, cDNA was synthesized using random hexamers (Superscript IV, Invitrogen), and PCR was performed with the following primers. *trm* mutant transcript analysis: forward 5’-GCATTGCATCTAA-AGGGCAGACGAA-3’ reverse 5’-TAAGGGTCCTGAGACTGCGGATAGA- 3’. *tfap2a* MO-splice transcript analysis: forward 5’-CCCTATCCATGGAATACCTCACTC-3’ reverse 5’-GATTACA-GTTTGGTCTGGGATGTGA-3’. *tfap2b* MO-splice transcript analysis: forward 5’-AGTGC-CTGAACGCGTCTCTGCTTGGT-3’ reverse 5’-TGACATTCGCTGCCTTGCGTCTCC-3’. For *tfap2a* MO and *tfap2b* MO transcript analysis, bands were gel-extracted, purified, and sequenced. For *trm* mutant transcript analysis, bands were gel-extracted, purified, and cloned into the pGemTEasy vector (Promega), and minipreps were sequenced.

### Alcian blue staining and o-dianisidine staining

Alcian blue cartilage staining was performed as previously described (Neuhauss et al., 1996). In brief, larvae were fixed at 4 dpf for 2 hours at RT in 4% PFA. Larvae were bleached for 1 hour in 10% KOH, 30% H_2_O_2_, 20% Tween diluted in distilled water. Samples were digested with proteinase K (10 mg/mL) diluted to 1X for 20 minutes. Samples were stained in 0.1% Alcian Blue (Sigma) dissolved in 70% ethanol / 5% concentrated HCl overnight, shaking at RT in glass vials. Larvae were destained using acidic ethanol for 4 hours, dehydrated by an ethanol series, and stored in glycerol. O-Dianisidine staining was performed as described on 4 dpf larvae to visualize blood and vasculature (Wingert et al., 2004).

### Acridine orange assay

Acridine orange (AO; Sigma A6014; 100 X) staining was performed on WT and *trm* mutants to analyze cell death (Kroeger et al., 2017; Westerfield, 193). In brief, a 50 X AO stock solution (250 µg/ml) was made. At 24 hpf, embryos were incubated in 1:50 AO solution (made from 50 X stock) diluted in 0.003% PTU/E3 media protected from light for 1 hour. Embryos were then washed three times with 0.003% PTU/E3, and then imaged with a dissecting microscope under the GFP filter in 2% methylcellulose/0.02% tricaine.

### Dextran clearance assay

To assess kidney function in WT and *trm* mutants, clearance assays using fluorescent 40 kDa dextran-fluorescein (FITC) (Invitrogen) were completed. Embryos were treated with 0.003% PTU at 24 hpf. At 2 dpf, embryos were anesthetized with 0.02% tricaine and dextran-FITC was injected into circulation. Live fluorescent imaging was performed 1 hour after injection and 24 hours after injection. Embryos were live-imaged with a dissecting microscope under the GFP filter in methylcellulose/0.02% tricaine.

## Acknowledgements

We thank Bruce Riley for sharing the inducible *hs:tfap2a* zebrafish transgenic with us. We thank the staffs of the Department of Biological Sciences and the Center for Zebrafish Research at the University of Notre Dame for their dedication and care of our zebrafish aquarium. Finally, we thank the members of our lab for their support, discussions, and insights about this work.

## Author Contributions

BEC, GFG, KHC, EGC, IL and RAW performed experiments. BEC, GFG, KHC, EGC, WG and RAW analyzed the results. BEC and RAW wrote and revised the paper.

## Data Availability

All data related to the present study is provided within the figures and supplementary information.

## Supplemental Figures & Figure Legends

**S1 Figure:**
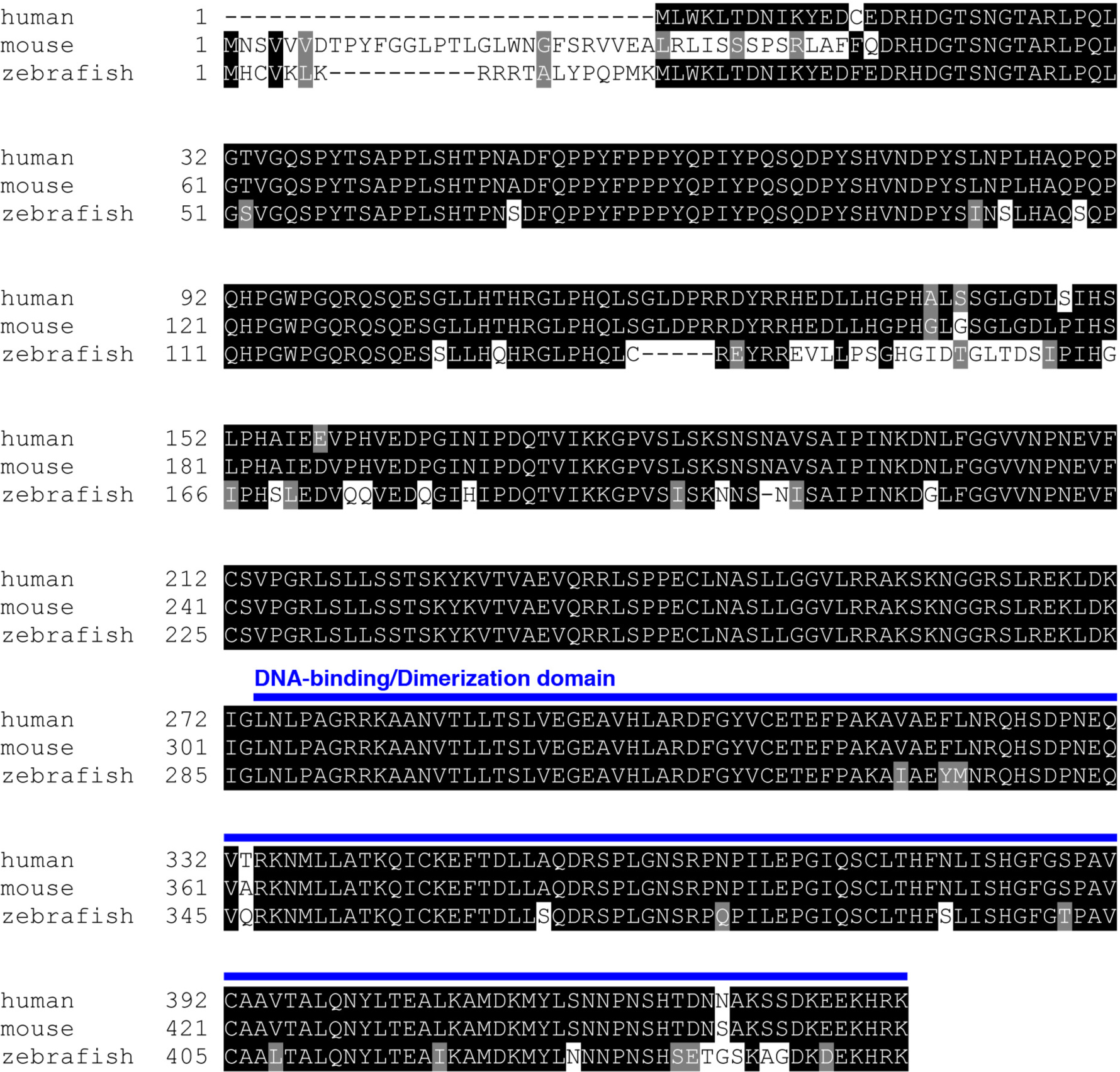
Tfap2a amino acid sequence is highly conserved across vertebrate species. Depicts amino acid alignment of human, mouse, and zebrafish Tfap2a generated by T-coffee and Boxshade online tools. DNA-binding and dimerization motifs (blue line) display a high degree of sequence similarly (>90 percent). Black boxes mark conserved residues.

**S2 Figure:**
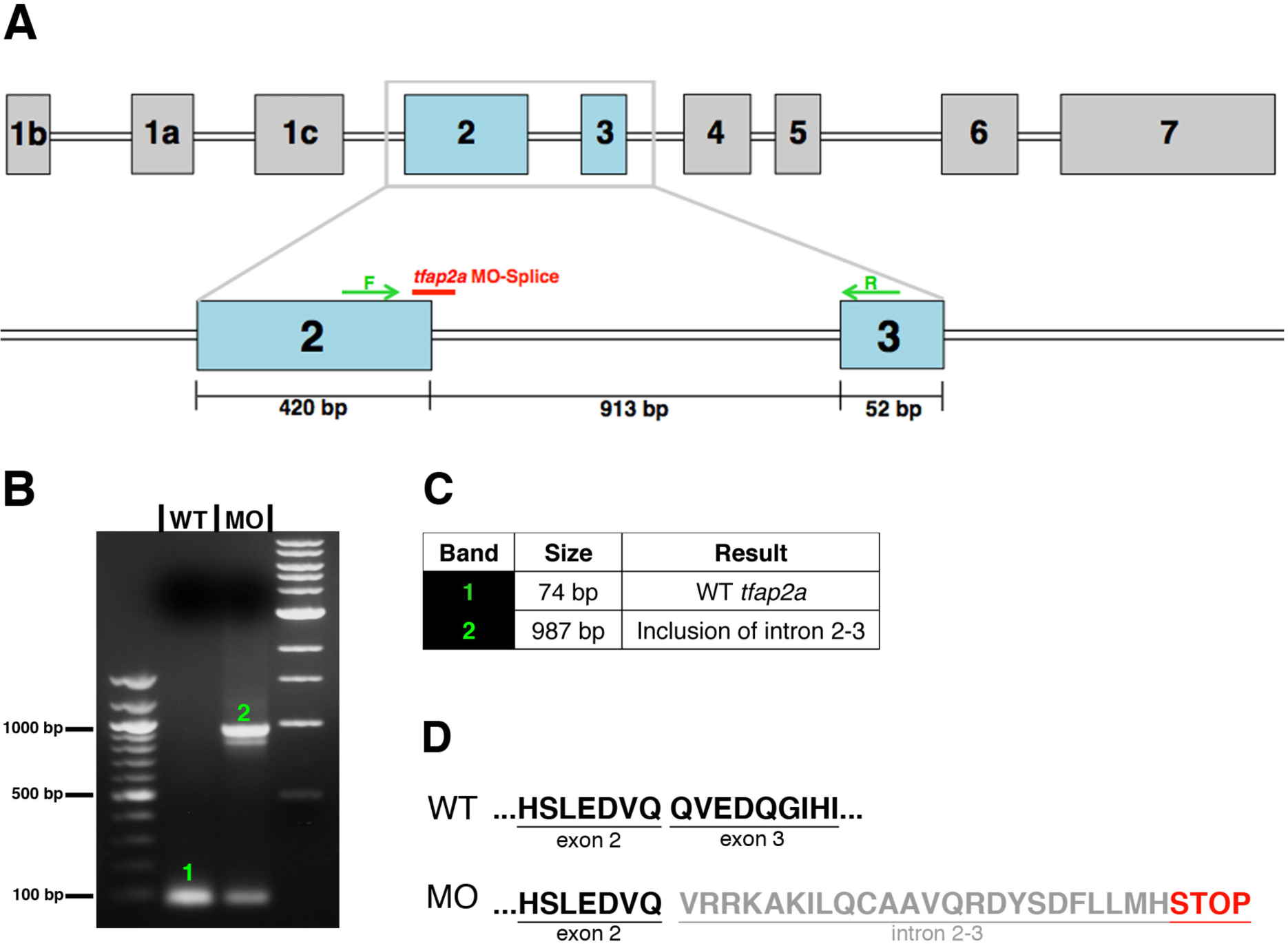
*tfap2a* MO splice efficacy verification through RT-PCR analysis. **A.** *tfap2a* exon map, indicating *tfap2a* MO-splice (red) targets the 3’ end of exon 2. Forward and reverse primers used for RT-PCR analysis are situated within exon 2 and exon 3 (green arrows). **B.** Image of RT-PCR gel reveals presence of a larger product size in the morphant lane, indicating disrupted splicing. 1 = WT band, 2 = morphant band. **C.** Table presenting the sequencing results of each band. **D.** WT and *tfap2a* morphant amino acid sequences. Inclusion of intron 2-3 results in premature stop codon (red), and a predicted truncated protein in *tfap2a* morphants.

**S3 Figure:**
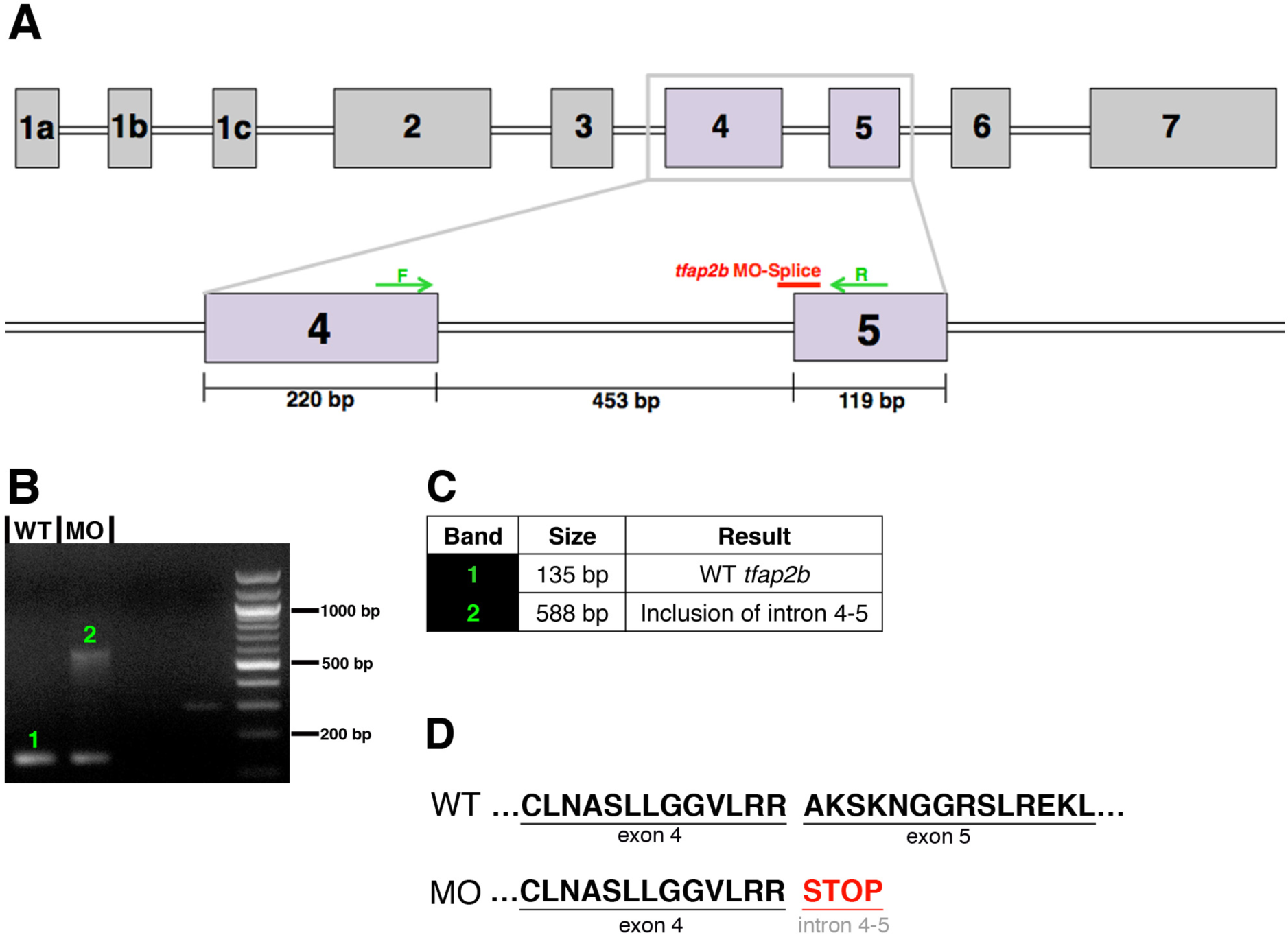
*tfap2b* MO splice efficacy verification through RT-PCR analysis. A. *tfap2b* exon map, indicating *tfap2b* MO-splice (red) targets the 3’ end of exon 4. Forward and reverse primers used for RT-PCR analysis are situated within exon 4 and exon 5 (green arrows). B. Image of RT-PCR gel reveals presence of a larger product size in the morphant lane, indicating disrupted splicing. 1 = WT band, 2 = morphant band. **C.** Table presenting the sequencing results of each band. **D.** WT and *tfap2b* morphant amino acid sequences. Inclusion of intron 4-5 results in premature stop codon (red), and a predicted truncated protein in *tfap2b* morphants.

**S4 Figure:**
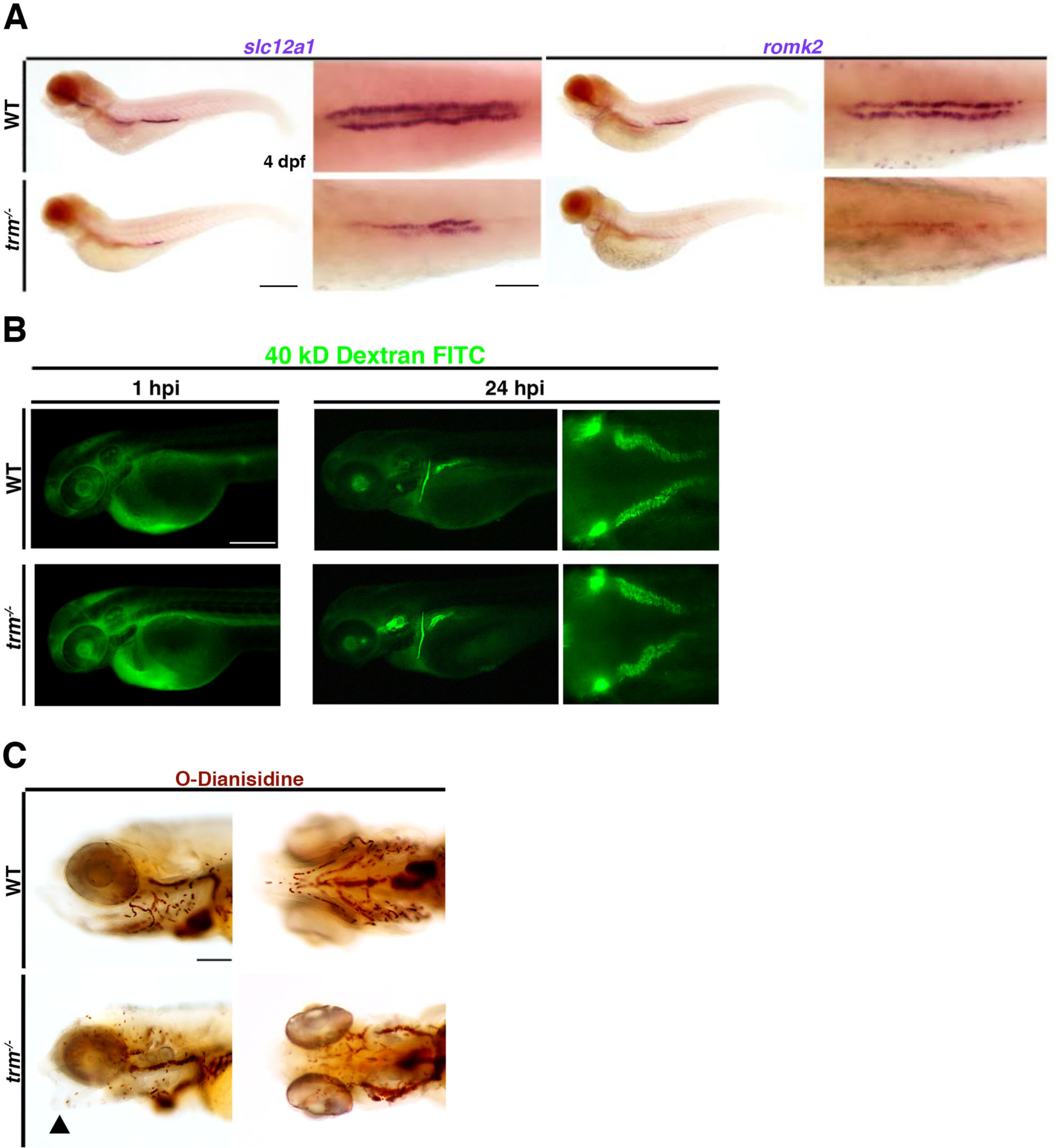
*trm*^-/-^ mutant embryos exhibit normal filtration and proximal tubule fluid uptake at 3 dpf. **A.** WISH analysis of DE markers (*slc12a1* and *romk2*) in WT and *trm*^-/-^ mutants at 4 dpf. Scale bars = 200 µm, 50 µm. **B.** Kidney function assay was performed by injecting 40 kD Dextran FITC into the circulation of 2 dpf WT and *trm*^-/-^ larvae. Images were collected 1-hour post injection and 24-hours post injection. Right panel: nephron tubules labeled with green fluorescence indicate endocytosis of Dextran. Scale bar = 150 µm. **C.** O-dianisidine staining of craniofacial vasculature in WT and *trm*^-/-^ at 4 dpf. Abnormal cartilage is annotated by black arrowhead. Scale bar = 100 µm.

**S5 Figure:**
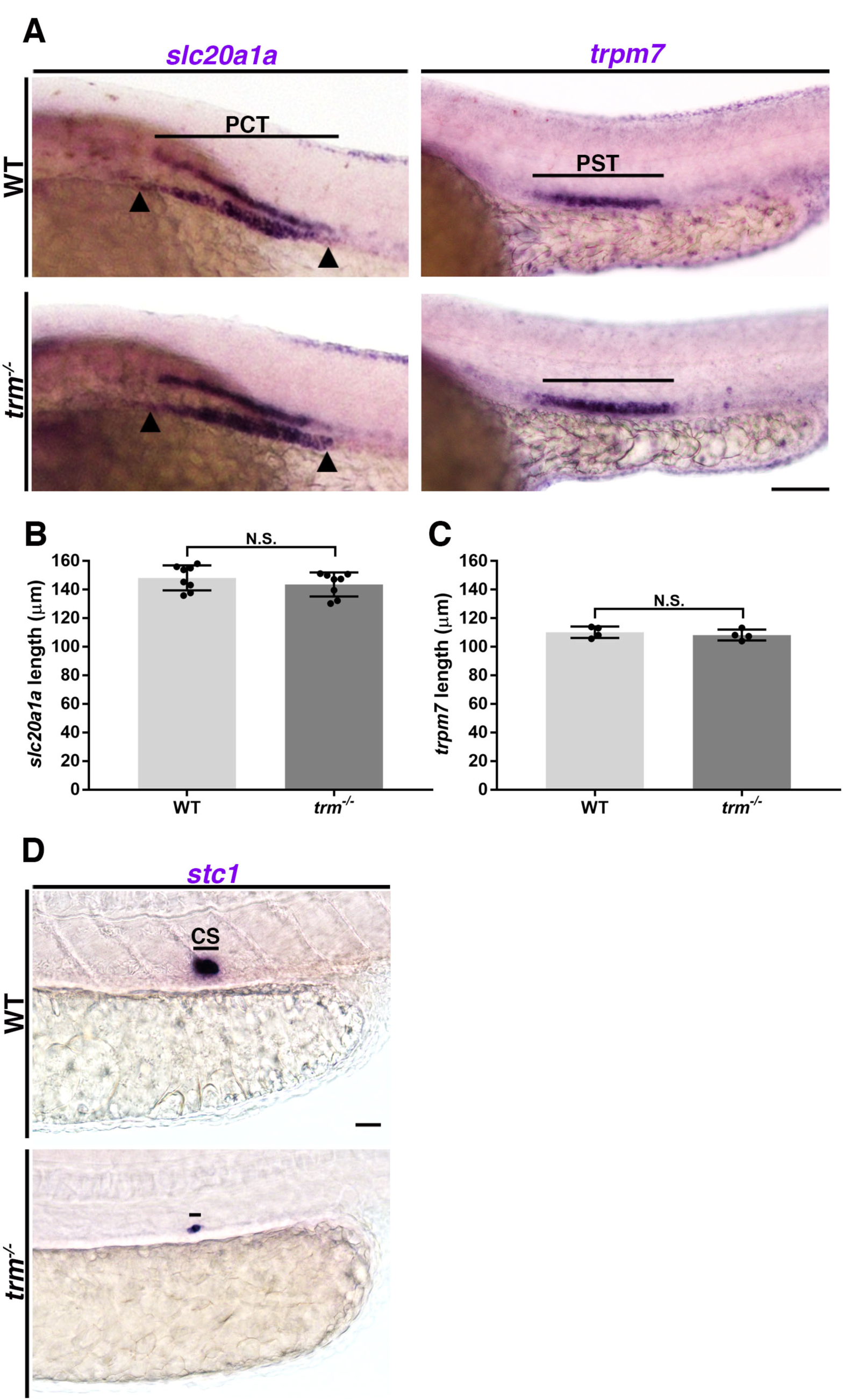
*trm*^-/-^ mutant embryos exhibit normal proximal nephron segment pattern formation but have abrogated corpuscle of Stannius formation. **A.** WISH to evaluate proximal convoluted tubule (*slc20a1a*) and proximal straight tubule (*trpm7*) in WT and *trm*^-/-^ at 24 hpf. Black arrowheads indicate the start and end of *slc20a1a* expression. Scale bar = 50µm. **B.** Quantification of absolute lengths of *slc20a1a* expression. **C.** Quantification of absolute lengths of *trpm7* expression. n = 3 for each control and test group. Measurements were compared by unpaired t-test. Data are represented as ± SD. N.S. = not significant. **D.** WISH to assess corpuscle of Stannius (*stc1*) development in WT and *trm*^-/-^ at 48 hpf. Black line indicates *stc1* expression domain. Scale bar = 20 µm.

## Key Abbreviations

(BOFS): branchio-oculo-facial syndrome
(CS): corpuscle of Stannius
(DE): distal early 27
(DL): distal late
(FISH): fluorescent whole mount *in situ* hybridization
(IF): immunofluorescence
(*irx1a*): *iroquois homeobox 1a*
(*irx3b*): *iroquois homeobox 3b*
(hpf): hours post fertilization
(MET): mesenchymal to epithelial transition
(MO): morpholino oligonucleotide
(PCT): proximal convoluted tubule
(PST): proximal straight tubule
(ss): somite stage
(TAL): thick ascending limb
(*tfap2a*): *transcription factor AP-2 alpha*
(*tfap2b*): *transcription factor AP-2 beta*
(WISH): whole mount *in situ* hybridization
(WT): wild-type

